# Higher ploidy coincides with inferior performance and no difference in stress tolerance in reed

**DOI:** 10.1101/2025.03.20.644296

**Authors:** Kerstin Haldan, Kristina Kuprina, Clemens Düsterhöft, Franziska Eller, Anke Fiehn, Aron Garthen, Nils Krauß, Martin Schnittler, Manuela Bog, Jürgen Kreyling

## Abstract

(1) Climate change leads to more extreme weather events. Therefore, a high stress tolerance is becoming more critical for plants, with higher ploidy being reported to lead to higher stress tolerance. *Phragmites australis* (*P. australis*) is a target species for paludiculture, i.e. the wet use of peatlands, and well known for its many ploidy levels.
(2) We expected octoploid genotypes of *P. australis* to outperform tetraploid ones in a 15-month mesocosm experiment including a gradient of 0 to 100 days of drought. We used pairs of genotypes differing in ploidy from three different geographic regions.
(3) Increasing drought length led to a decrease in growth, biomass, morphological and ecophysiological traits in both ploidy levels, but 4× outperformed 8× in almost all traits under constant water supply (e.g., 2.5-fold more biomass production) and up to moderate drought (about 50 days). Under severe and prolonged drought, both ploidy levels performed equally poorly.
(4) Our study suggests that higher ploidy levels do not necessarily outperform lower ploidy levels of *P. australis* under stressful conditions. In this species, ploidy alone may not explain performance, but the genotype can be as or more important than ploidy.

## Introduction

Polyploidy, i.e. possessing more than two complete sets of chromosomes, is a common occurrence in plants (Comai, 2005; One Thousand Plant Transcriptomes Initiative, 2019). Many studies, across various plant species and environmental stressors, suggest that polyploid plants show an increased stress tolerance compared to their diploid relatives (Tossi *et al*., 2022). As climate change leads to more frequent and intense extreme events, such as droughts, floods and heat waves (IPCC, 2023), an increased stress tolerance of plants is critical and especially relevant in agriculturally used species (Xiong *et al*., 2006; Bailey-Serres *et al*., 2012; Dong *et al*., 2020; Wójcik *et al*., 2022; Abdolinejad & Shekafandeh, 2022). Exposed to environmental stress, polyploids, compared to their diploid counterparts, show higher photosynthesis or less reduction in photosynthesis or other physiological parameters, better survival, growth and germination, a stronger antioxidative response, stronger hormonal stress response, less stress markers and a different gene expression and transcription (Deng *et al*., 2012; Xue *et al*., 2015; Zhu *et al*., 2018; Akinroluyo *et al*., 2019; Lu *et al*., 2020; Talebi *et al*., 2021; Wójcik *et al*., 2022; Abdolinejad & Shekafandeh, 2022; Jin *et al*., 2022; Wang *et al*., 2022; Tossi *et al*., 2022). Only about 10% of studies mention a negative or no effect of polyploidy on stress tolerance (Chandra & Dubey, 2009; Oustric *et al*., 2019; Dwivedi *et al*., 2022; Tossi *et al*., 2022).

*Phragmites australis* (CAV.) STEUD. (*P. australis*), the common reed, is a globally distributed hygrophyte known for its many ploidy levels (3×, 4×, 6×, 7×, 8×, 10×, 11×, 12×) of which tetraploids (4×) and octoploids (8×) are most common (Gorenflot, 1976; Clevering & Lissner, 1999; Packer *et al*., 2017). In *P. australis*, polyploidy is not generally related to habitus (e.g. plant size) or ecotype (Gorenflot, 1976; Clevering & Lissner, 1999; Hansen *et al*., 2007). Only few differences in morphology (larger cell sizes, longer guard cells and lower stomatal density in higher ploidy levels; lower growth density and taller or smaller growth of 8× compared to 4×), seasonality (longer growing season of 8× compared to 4×) and phytochemistry (better chemical defence of 4× compared to 8×) have been reported between ploidy levels (Pauca-Comanescu *et al*., 1999; Hansen *et al*., 2007; Achenbach *et al*., 2012; Meyerson *et al*., 2016).

Previous studies provide contrasting results on whether *P. australis* ploidy levels react differently to environmental stress. On the one hand, 8× were found to be more affected by salt stress than 4×, on the other hand, salt stress reduced differences between ploidy levels or no effect of ploidy level was found (Hanganu *et al*., 1999; Pauca-Comanescu *et al*., 1999; Achenbach *et al*., 2013). The response of different ploidy levels of *P. australis* to other stressors than salt stress has hardly been studied so far.

Moreover, many of these differences were observed while comparing single genotypes, without replication within ploidy levels, despite the fact that genotype and origin can have a major influence on the phenotype in *P. australis*. A notable exception are Meyerson *et al*. (2016) who used well over 100 genotypes in their study. Consequently, further studies found no differences in guard cell size, stomatal density, biomass allocation and physiological traits between ploidy levels or differences in these traits could be attributed to subspecies or genotype instead of ploidy level (Pauca-Comanescu *et al*., 1999; Saltonstall *et al*., 2007; Hansen *et al*., 2007; Achenbach *et al*., 2012). Ultimately, studies on different ploidy levels of *P. australis*, which exclude an influence of genotype and origin, are very rare and insufficient to give a coherent picture.

*P. australis* has always been used in numerous ways. Nowadays, it is gaining importance again as a suitable crop for paludiculture, a form of agriculture in which drained fen peatlands are rewetted and subsequently cultivated for climate protection (Wichtmann & Joosten, 2007). Globally, about 12% of peatlands have been destroyed to the point where they no longer produce peat, but instead their peat is being decomposed, and a further 500,000 ha of peat-forming peatlands are destroyed annually by human activities (UNEP, 2022). Degraded peatlands emit ca. 2,000 megatons of CO2-equivalents annually, which is about 4% of man-made greenhouse gas emissions, and drainage terminates important ecosystem services, such as carbon sequestration, water retention/regulation and harbouring unique biodiversity. Rapid carbon emission reduction and a stop to further degradation can be achieved by rewetting drained peatlands (Evans *et al*., 2021, Günther *et al*., 2020).

Drought is a major environmental stress. Hydrology of natural wetlands can be highly variable and even greater water table amplitudes can occur in rewetted peatlands (Kreyling *et al*., 2021). Climate change will likely lead to more frequent and intense droughts (Trenberth *et al*., 2014; IPCC, 2023) which are predicted to result in drastic losses in crop yield (Li *et al*., 2009). Any information on whether *P. australis* of different ploidy show different resistance to environmental stresses could prove very helpful for future cultivation in paludiculture.

While *P. australis* commonly occurs in wet environments and is adapted to flooding, it can tolerate dry conditions and prolonged drought (Gorenflot, 1976; Pagter *et al*., 2005). The overall negative effect of drought on ecophysiological processes and growth of *P. australis* is similar to other wetland species (Touchette *et al*., 2010; Didiano *et al*., 2018; da Cunha Cruz *et al*., 2019; Huang *et al*., 2022). In *P. australis*, even mild water stress can affect some plant traits, e.g. lead to reduced leaf production and growth and thus reduced leaf area per plant (Pagter *et al*., 2005). When water stress becomes more severe, leaves roll up, wilt and ultimately die off and are shed (Pagter *et al*., 2005; Saltmarsh *et al*., 2006; Naumann *et al*., 2007). On a whole-plant level, growth declines, then stops completely and ultimately leaves and stems die off resulting in a decrease in the percentage of living biomass and reduction of total standing biomass under drought stress (Saltmarsh *et al*., 2006; Liu *et al*., 2018). Other plant traits of *P. australis* may be favoured by short drought, indicating that permanent flooding is not optimal for the species. In the first six days of drainage, CO2 assimilation rate and shoot elongation rate were higher than under flooded conditions and only decreased under prolonged drainage (Saltmarsh *et al*., 2006). Likewise, photosynthesis remained at a high level until very low water availability, indicating that leaves stayed physiologically active until they suffered severe stress (Pagter *et al*., 2005; Saltmarsh *et al*., 2006). Only at a low water availability did CO2 assimilation rate, intercellular CO2 concentration, transpiration rate and chlorophyll fluorescence decline, but no effect of drought on chlorophyll content was observed (Pagter *et al*., 2005; Saltmarsh *et al*., 2006). Furthermore, *P. australis* can counter drought stress with mechanisms such as osmotic adjustment and increased water use efficiency (Pagter *et al*., 2005; Liu *et al*., 2018).

However, the response of *P. australis* to drought stress has mainly been studied in small-scale and short-term experiments (Pagter *et al*., 2005; Saltmarsh *et al*., 2006; Naumann *et al*., 2007) and the response of different ploidy levels of *P. australis* to drought has not been studied at all.

In the current study we present novel evidence of changes in growth, productivity and ecophysiology of tetraploid (4×) and octoploid (8×) *P. australis* genotypes in response to drought. To this end, we cultivated pairs of 4× and 8× *P. australis* genotypes from three distinct geographic origins in a mesocosm experiment and subjected them to a gradient of 0 to 100 days of drainage. We hypothesized that *P. australis* with the higher ploidy level (8×) can cope better with drought than *P. australis* with the lower ploidy level (4×). 8× genotypes will cease growth and their biomass will die off at more severe stress than 4× genotypes, leading to more biomass at harvest of 8× compared to 4×.

## Material and Methods

### Experimental setup

We investigated growth, biomass, morphology and photosynthesis of tetraploid (4×) and octoploid (8×) *Phragmites australis* (CAV.) STEUD. (*P. australis*) from three distinct geographic origins in response to drought.

Six *P. australis* genotypes were obtained from the common garden collection at Aarhus University (56°13’43.5’ N 10°07’37.4’ E) in November 2019. The plants used in this study were propagated meristematically from rhizome buds from November 2019 to June 2020. To control for any influence of geographic origin, we selected three pairs of genotypes. Each pair consisted of a 4× and an 8× genotype from one origin (Table 1), i.e. Lake Fertő in Hungary (genotypes Hu 4× and Hu 8×), Lake Razim in Romania (Ro 4× and Ro 8×) and Sakhalin Island in Russia (Ru 4× and Ru 8×). Ploidy levels of all genotypes were confirmed using flow cytometry.

**Table 1:** The six *Phragmites australis* genotypes used in this study, consisting of three tetraploid (4×) / octoploid (8×) pairs from three distinct geographic origins, were obtained from the common garden collection at Aarhus University.

| **Label** | **Ploidy** | **Country** | **Description of locality** | **Approximate coordinates** |
| --- | --- | --- | --- | --- |
| Hu 4x | 4x | Hungary | Lake Fertő; nearest Place Kapuvar | 47°36'00.0'' N<br>17°001'36.0'' E |
| Hu 8x | 8x | Hungary | Lake Fertő; nearest Place Kapuvar | 47°36'00.0'' N<br>17°001'36.0'' E |
| Ro 4x | 4x | Romania | Lake Razim | 45°00'00.0'' N<br>29°013'00.0'' E |
| Ro 8x | 8x | Romania | Lake Razim | 45°00'00.0'' N<br>29°013'00.0'' E |
| Ru 4x | 4x | Russia | Sakhalin Island; Novikovo<br>(Korsakovsky District) | 46°37'12.0'' N<br>142°46'048.0'' E |
| Ru 8x | 8x | Russia | Sakhalin Island; Yuzhno-Sakhalinsk<br>B.G. | 46°57'08.0'' N<br>142°44'026.0'' E |

The experiment was set up as a mesocosm gradient experiment (Kreyling et al., 2018) at the Arboretum of Greifswald University (54°05’31.4’ N 13°24’36.5’ E) with 11 treatment levels, ranging from 0 to 100 days of drought in steps of ten days.

Plants were grown in plastic tubes (diameter 20 cm, height 60 cm) filled with a substrate of two parts white peat (pH 6, Rostocker Humus & Erden GmbH, Rostock, Germany) and one part sand. Tubes were sealed at the bottom with water-permeable root fleece (polypropylene, 150 g m^-2^). Two clones per tube were planted on 8^th^ June 2020 and received 0.744 g of slow-release fertilizer (Basacote Plus 6M, 16 N + 8 P + 12 K (+ 2 Mg + 5 S) + trace elements, Compo Expert, Münster, Germany). The two clones planted per tube were not distinguishable from each other over the course of the experiment and were therefore treated – and are referred to in this paper – as one plant.

Per treatment level, one tube of each genotype was placed inside a 1000L Container (1 x 1 x 1 m) filled with communal tap water (Table S1). Tubes were arranged randomly inside the container and surrounded by 12 tubes with further *P. australis* plants, on which response parameters were not quantified, to avoid edge effects. Containers, each representing a treatment level, were arranged in a random order and without shading each other.

Water level was kept constant at soil surface. Three soil moisture sensors (TMS-4, TOMST s.r.o., Prague, Czech Republic) were installed per treatment level (see Supporting Information Methods S1 for calculation of volumetric soil moisture). To keep out rain, containers were covered with roofs made of greenhouse foil (Lumisol PGH-UV5 /UV B open, UV-B permeable, light permeability 88%, Poppen Gewächhausbau, Edewecht-Jeddeloh, Germany) from 11^th^ August 2020 to 18^th^ September 2020 and from 18^th^ May 2021 to 7^th^ September 2021. To monitor ambient conditions throughout the experiment, we used an air temperature and humidity sensor (VP-3 Humidity/Temp, Meter Group AG, Munich, Germany), one photon flux sensor (QSO-S PAR photon flux, Meter Group AG, Munich, Germany) outside the roofs and one photon flux sensor under a roof at approx. 50 cm above soil surface, connected to a data logger (Decagon Em 50 Data Logger, Meter Group AG, Munich, Germany).

In 2020, plants already received a drought treatment which, however, showed no effect on plant performance, probably due to water stress not being reached (pF 2.2 to 2.8), because of drought onset too late in the season (August) and too mild treatments (0 to 40 days) combined with plants not yet requiring much water as they still were small. Details, including a slight positive effect of previous drought on height before the start of the main treatment for 8×, can be found in Supporting Information Methods S2.

By 11^th^ April 2021, water level was raised to soil surface and kept there aside from the drought treatments. Plants were fertilized weekly for 7 weeks from April 9^th^ onwards with 15 g of Hakaphos Blau (15N+10P+15K+2Mg+trace elements; Compo Expert, Münster, Germany), and, from the second fertilization onwards, additionally with 0.5 g iron chelate (YaraTera® TENSO® IRON 58, 6% water soluble iron, 4% HBED iron chelate, 1.8% EDDHMA iron chelate, YARA GmbH & Co. KG, Dülmen, Germany) dissolved in 0.25 L tap water per tube.

In 2021, plants were subjected to a gradient of drought with 11 treatment levels, ranging from 0 (no drought treatment) to 100 days drought length. Starting on 20^th^ May 2021, water was drained from another container every 10 days so that the drought ended at the same date for all levels.

### Response parameters

Maximum plant height (indicating plant growth) was measured weekly from 9^th^ April to 18^th^ August 2021 and again on 28^th^ August before harvest. Maximum plant height per tube was determined by gently holding the stem and leaves upright and measuring to the highest green part of the plant. Stems were counted only including alive stems (showing some green on the stem) and stems higher than 3 cm.

Photosynthesis was determined by leaf gas exchange using the LCi-SD Leaf Chamber Analysis System (ADC BioScientific Limited, Hoddesdon, Herts, England) with external PLU5 LED light unit (given PAR, PAR Qleaf (g) = 1577 µmol m^-2^ s^-1^, PAR Qgiven = 1715 µmol m^-2^ s^-1^) fitted with the narrow leaf chamber without radiation shield (area: 5.80 cm^2^, Hfac: 0.168, rb: 0.30, Tlmtd: measured, Trw=0.92, uset = 200 µmol s^-1^). Photosynthesis was measured on four days (25^th^ August to 28^th^ August 2021) at the end of the drought treatment. To avoid a daytime influence on photosynthesis readings, each plant was measured once in the morning, once around noon and once in the afternoon. On each plant, photosynthesis was measured on two leaves. On leaves too small for the measurement chamber (5.8 cm x 1 cm), leaf width at each end of the chamber (*a*, *c*) and greatest leaf width in the middle of the chamber (*b*) were measured. Photosynthesis was corrected for photosynthetically active leaf area in the chamber using formula 1:

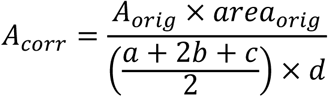

with *A*_*corr*_ being the corrected photosynthesis, *A_orig_* the originally measured photosynthesis, *area_orig_* the chamber area (5.8 cm^2^), *a*, *b* and *c* the leaf width measurements and *d* half the chamber length (2.9 cm). From all measurements on a plant (3 times x 2 leaves) a mean value per genotype and treatment level was calculated.

Aboveground biomass harvest took place from 28^th^ August to 3^rd^ September 2021, starting directly after the end of the drought treatments. All stems were cut at soil surface. On the ten longest stems per tube, total stem length (to the tip of the upper leaves gently extended along the stem) and stem diameter (mean of two orthogonal measurements at the base of the stem) were measured and all leaves on the stem counted and categorised as alive (leaf at least in part green) or dead (leaf without green parts). Then, all stems per tube were counted and categorised as alive, if a stem had at least one alive leaf, or dead, if a stem had no alive leaves. Biomass was divided into an alive and a dead fraction per plant, with all stems categorised as alive making up the alive fraction and all stems categorised as dead making up the dead fraction. To determine photosynthetically active leaf area, the third fully developed leaf of the three longest stems per genotype and level was harvested and stored in a sealed plastic bag with a moist paper towel. Leaves were dabbed dry before scanning with a flatbed scanner (400 dpi, reflected light, film type positive film, document type film area guide; Epson Perfection V800 Photo, EPSON Deutschland GmbH, Meerbusch, Germany). Photosynthetically active leaf area was analysed using ImageJ (Version 1.53k, Schneider *et al*. (2012)), colouring photosynthetically active leaf area using color threshold (thresholding method default, threshold color red, color space HSB, Hue 42 – 170, saturation 0 – 255, Brightness 0 – 255), then measuring it using ImageJ. A mean value of the three leaves per genotype and level was calculated.

Biomass was dried at 60°C until constant dry weight (after 120 h) and weighed (Kern PCB, d = 0.01 g, KERN & SOHN GmbH, Balingen-Frommern, Germany).

Belowground biomass harvest took place on 6^th^ and 7^th^ September 2021, when root systems were washed coarsely, placed into plastic bags and stored at 4°C until further processing. Root systems were carefully washed free of substrate and separated into roots and rhizome. Roots and rhizomes were dried at 60°C until constant dry weight (72 h) and weighed (Kern PCB, d = 0.01 g, KERN & SOHN GmbH, Balingen-Frommern, Germany). Root systems too big to be processed completely (> 1 kg) were divided into two parts, weighed (Kern CH, d = 0.1 kg, KERN & SOHN GmbH, Balingen-Frommern, Germany) and only one part was processed. Root and rhizome weight were extrapolated to the total weight of the root system.

For calculation of biomass allocation see Supporting Information Methods S3.

### Data analysis

To explore the shape of plant response to drought, this study was designed as a gradient experiment (Kreyling *et al*., 2018). We expected non-linear response patterns along the drought gradient and used generalized additive modelling (GAM) to unravel these patterns. Each dependent variable was modelled as a smooth function over drought length separately for each ploidy level (4× or 8×, Table S2). Ploidy was added to the model as a factor by-variable in the smooth term and as main effect. A thin plate regression spline was used as the smooth function (bs = “tp”). Restricted maximum likelihood was used as the smoothing parameter estimation method (method = “REML”). Number of basis functions (k) was restricted only if the original graph appeared unusually wiggly or had multiple maxima. The effect of drought length on the dependent variable, separately for each ploidy level, was considered significant (α = 0.05) if the p-value of the respective smooth term was below 0.05. Significant difference of the two ploidy levels along the drought gradient was evaluated visually: The two ploidy levels were considered significantly different (α = 0.05) if their 83% confidence intervals did not overlap (Austin & Hux, 2002; Payton *et al*., 2003; Knol *et al*., 2011). Model fit, namely model convergence, number of basis dimensions, normality of residuals and homogeneity of variance of residuals were checked using gam.check (Table S3).

All analyses were performed in R (Version 4.2.1; R Core Team, 2022) using the packages mgcv (Version 1.8-42, Wood, 2003) for GAM and ggplot2 (Version 3.3.6, Wickham, 2016) for plotting.

## Results

### Soil moisture

Drainage led to an immediate decline in volumetric soil moisture from around 50% to 25% followed by a slower drying down to below 5% (Figure 1). Containers that were drained first dried out more slowly than containers that were drained later. The permanent wilting point (PWP, 5.2%) was reached on average 24 days after drainage. Only levels with 0, 10 and 20 days drainage did not reach PWP.

**Figure 1:**
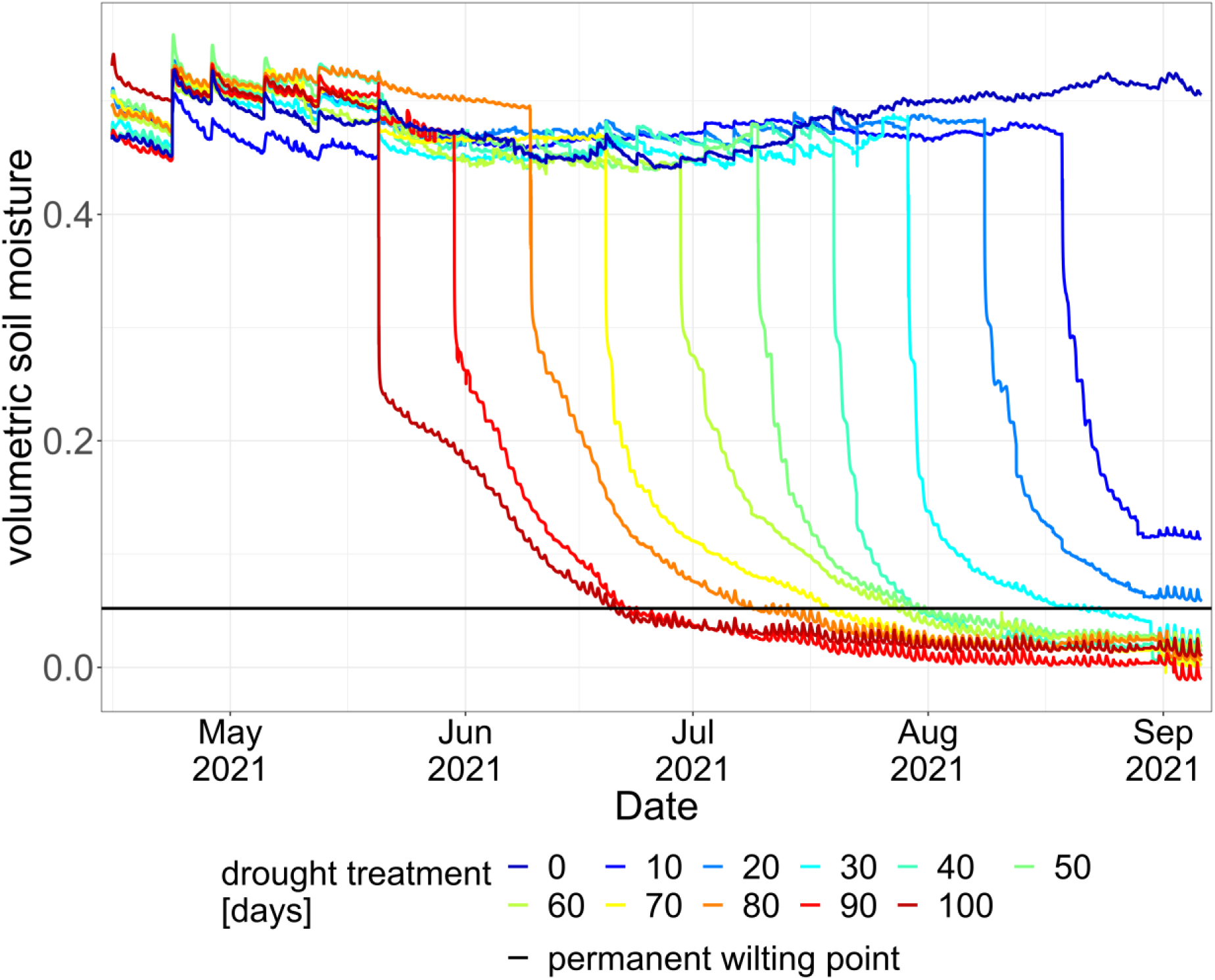
Volumetric soil moisture over the course of the experiment in all treatment levels from 0 (dark blue) to 100 days of drainage (dark red). Mean value of three soil moisture sensors per treatment level. The black line indicates the permanent wilting point (PWP, 5.2%).

### Growth

Increasing drought length led to a cessation of height growth and subsequent decline of living height towards the end of the treatment (Figure 2). In the no drought and 10 days drainage treatments, both 4× and 8× reached their tallest height at the end of the treatment (Table 2). In the other treatments, the tallest height was reached earlier and height declined significantly towards the end of the treatment. Height of 4× declined by 10%, 22%, 55%, 44%, 89% and 94% in the treatments with 20, 30, 40, 50, 60 and 90 days of drainage, respectively, and by 100% in the treatments with 70, 80 and 100 days drainage. Height of 8× declined by 10%, 16%, 21%, 52% and 64% in the treatments with 20, 30, 40, 50 and 60 days of drainage, respectively, and by 100% in the treatments with 70 and more days drainage. Time between onset of drought and cessation of height growth was longer for the longer drought treatments which started earlier in the growing season compared to a later onset of drought. As the growing season progressed, 4× grew significantly taller than 8× in treatments without or with short drought. An early onset of drought mostly prevented height differentiation between 4× and 8× resulting in a similar height of both ploidy levels throughout the duration of the experiment.

**Figure 2:**
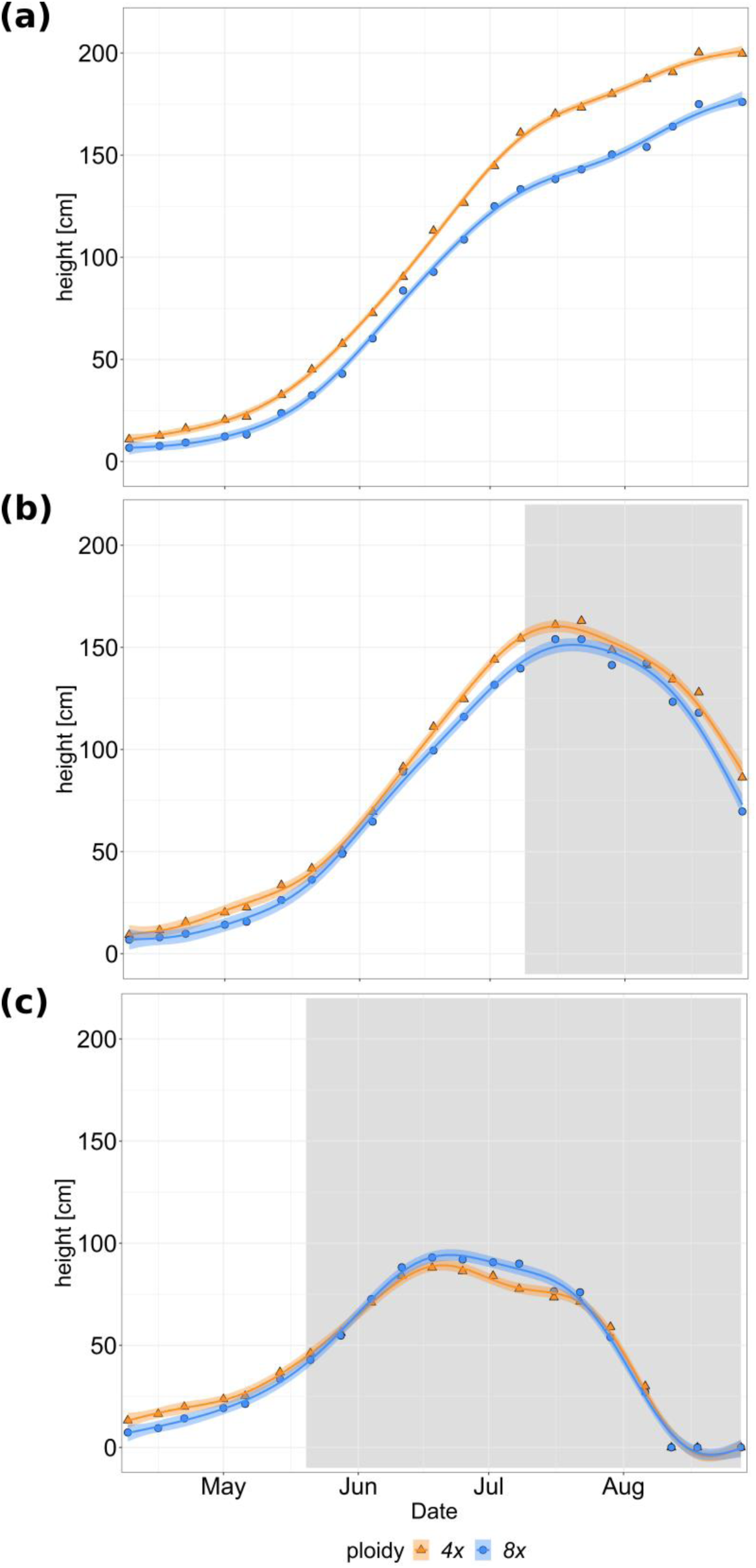
Height [cm] growth during the growing season 2021 of 4× (orange, triangles) and 8× (blue, circles) *Phragmites australis* genotypes in the treatments (a) no drought, (b) 50 days drainage and (c) 100 days drainage. Lines show the generalized additive model; shaded areas show the 83% confidence intervals. Gray shading in the background depicts the time period of drainage in the respective treatment

**Table 2:** Maximum and minimum fitted values of each measured parameter for tetraploid (4×) and octopoid (8×) *Phragmites australis* genotypes, obtained from the generalized additive models.

| Parameter | maximum<br>fitted<br>value 4x | maximum<br>fitted<br>value 8x | minimum<br>fitted<br>value 4x | minimum<br>fitted<br>value 8x |
| --- | --- | --- | --- | --- |
| **Growth** |  |  |  |  |
| Height [cm] | 206.4 * | 177.8 ** | - | - |
| Total number of stems | 95 ** | 64 † | - | - |
| **Biomass at harvest** |  |  |  |  |
| total biomass [g] | 414.54 | 141.54 | 33.35 | 24.47 |
| aboveground biomass [g] | 306.09 | 102.16 | 20.45 | 10.64 |
| belowground biomass [g] | 107.53 | 39.38 | 12.57 | 13.83 |
| **Morphology at harvest** |  |  |  |  |
| Stem length [cm] | 193.4 | 138.8 | 73.8 | 78.5 |
| Total number of stems | 103 | 67 | 51 | 44 |
| Alive number of stems | 10 | 11 | 1 | 1 |
| Mean stem diameter [mm] | 5.1 | 3.5 | 2.2 | 2.3 |
| Total leaves per stem | 11 | 13 | 6 | 7 |
| Alive leaves per stem | 10 | 11 | 1 | 1 |
| **Dead biomass fractions** |  |  |  |  |
| Fraction of dead aboveground<br>biomass in total aboveground biomass | 0.55 | 0.45 | 0 | 0 |
| Fraction of dead in total number of stems | 0.81 | 0.74 | 0.18 | 0.26 |
| Fraction of dead leaves in total number<br>of leaves per stem | 0.79 | 0.84 | 0.13 | 0.14 |
| **Biomass fractions** |  |  |  |  |
| Aboveground mass fraction | 0.75 | 0.74 | 0.61 | 0.57 |
| Belowground root mass fraction | 0.52 | 0.52 | 0.26 | 0.33 |
| **Photosynthetic rate** |  |  |  |  |
| Photosynthetic rate [$\mu\text{mol m}^{-2} \text{s}^{-1}$] | 11.47 | 13.36 | 1.33 | -0.53 |
| Photosynthetically active leaf area [$\text{cm}^2$] | 64.59 | 50.32 | -1.01‡ | -0.90‡ |

\* 10 days drought treatment, \*\* 0 days drought treatment, † 30 days drought treatment, ‡ these negative estimated values are ecologically impossible and should be interpreted as zero

Development of stem number followed the same general pattern as height, for details see Supporting Information Notes S1, Figure S1.

### Biomass and morphology at harvest

Biomass and morphological parameters at harvest of both 4× and 8× decreased significantly with increasing length of the drought treatment (see Figure 3a for total biomass, Figure 3b for height, Figure S2a for aboveground biomass and Figure S2b for belowground biomass, Figure S3a for total number of stems, Figure S3b for alive number of stems and Figure S3c for mean stem diameter, Figure S4a for total number of leaves per stem and Figure S4b for number of alive leaves per stem).

**Figure 3:**
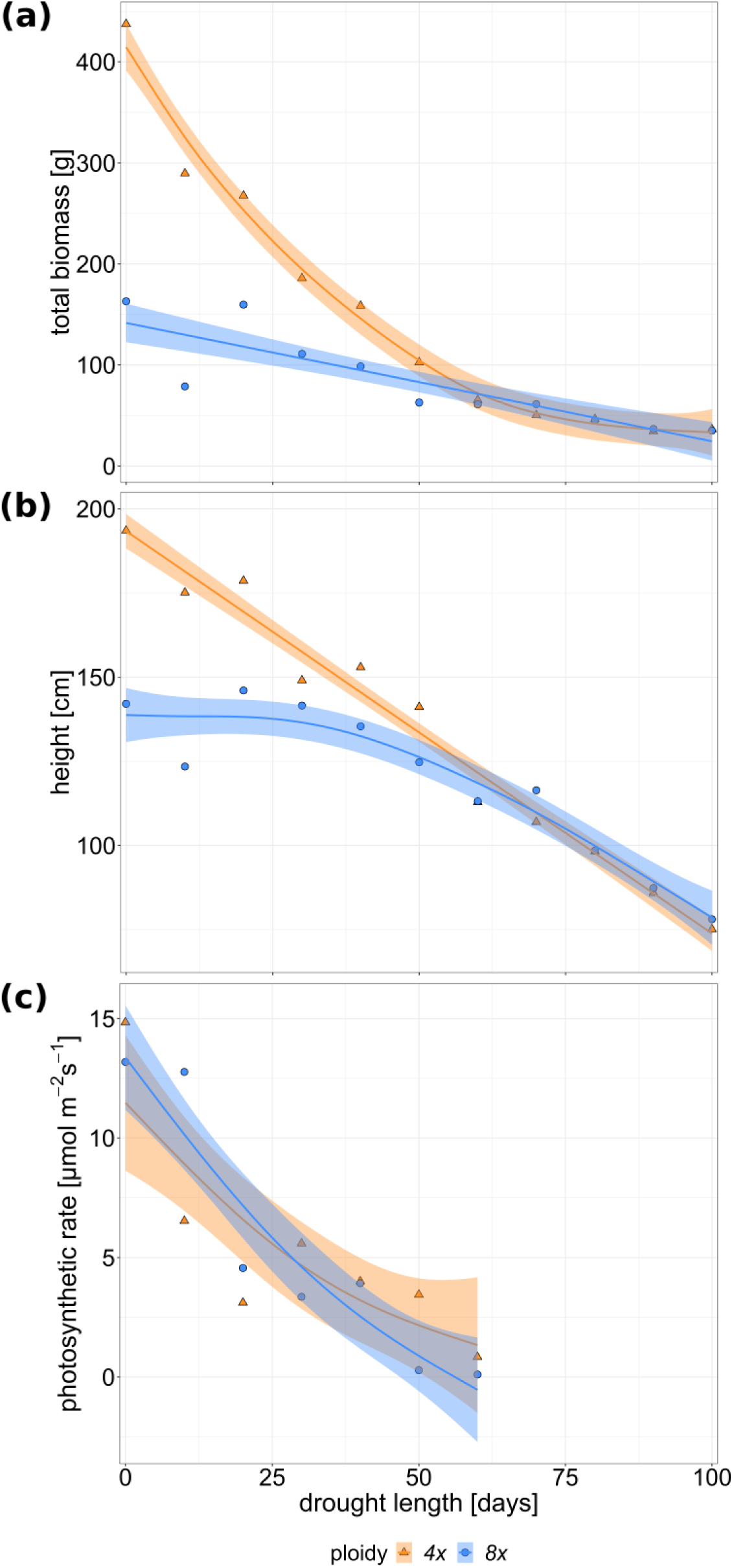
(a) Total biomass [g], (b) height [cm] at harvest and (c) photosynthetic rate [µmol m^-2^ s^-1^] of 4× (orange, triangles) and 8× (blue, circles) *Phragmites australis* genotypes along the drought length gradient. Height at harvest was measured as the mean length of the 10 longest stems of each genotype per level. Lines show the generalized additive model ((a) total biomass: adjusted R^2^ = 0.951, 4×: edf = 3.466, F-value = 76.57, p < 0.001, 8×: edf = 1, F-value = 29.23, p < 0.001; (b) height at harvest: adjusted R^2^ = 0.954, 4×: edf = 1, F-value = 321.28, p < 0.001, 8×: edf = 2.693, F-value = 28.96, p < 0.001; (c) photosynthetic rate: adjusted R^2^ = 0.755, 4×: edf = 1.586, F-value = 9.717, p < 0.05, 8×: edf = 1.42, F-value = 18.107, p < 0.01). Shaded areas show the 83% confidence intervals.

Without drought, 4× were about three times as productive (Figure 3a), and taller at harvest (Figure 3b), than 8× (Table 2). 4× remained significantly more productive and taller than 8× up to a treatment of about 40 to 60 days drainage, depending on the response parameter. Under the impact of even longer drought treatments, values of morphological measurements and biomass at harvest of 4× and 8× converged at low values (Figures 3, S2, S3, S4, Table 2). Notably, biomass of 4× decreased approximately exponentially, while biomass of 8× decreased linearly, meaning that especially under short drought, biomass of 4× decreased much stronger than that of 8×, so that almost the entire decrease occurred up to a treatment of 50 days drainage and flattened out during even longer drought. An exception from this trend were number of total and alive leaves per stem, of which 8× had more than 4× at up to about 60 days drainage (Figure S4a,b, Table 2).

Fraction of dead aboveground biomass in total aboveground biomass increased significantly with prolonged drought (Figure S5a; Table 2; see also Figure S5b for fraction of dead stems, Figure S5c for fraction of dead leaves per stem). 4× had a higher fraction of dead stems and a higher fraction of dead leaves per stem than 8× in some treatments (Table 2). Effect of drought length on biomass allocation is described in Supporting Material Notes S2.

### Photosynthetic rate

Photosynthetic rate of both ploidy levels decreased significantly with increasing drought length (Figure 3c, Table 2). Photosynthetically active leaf area of both ploidy levels decreased immediately and steeply even at short drought treatments, the decrease was significant over the drought gradient (Figure S4c, Table 2). Ploidy levels did not differ in photosynthetic rate nor photosynthetically active area per leaf along the gradient.

### Genotypes

For some parameters, the individual genotypes differed greatly, with the general patterns between 4× and 8× described above not being observable in all geographic origins of the ploidy pairs. In several biomass and morphological characteristics, 4× were more productive and bigger than 8× in the Hungarian and Romanian origin pairs; this difference was less pronounced (total biomass, belowground biomass), not existent (height and total number of stems at harvest, mean stem diameter) or even opposite (aboveground biomass, number of alive stems at harvest) in the Russian origin pair. For instance, Hungarian and Romanian 4× were significantly higher at harvest than their 8× counterparts, at treatments of up to about 30 days drainage (Romanian genotypes) and 80 days drainage (Hungarian genotypes) while the Russian 4× and 8× did not differ significantly in height along the entire drought gradient (Figure 4).

**Figure 4:**
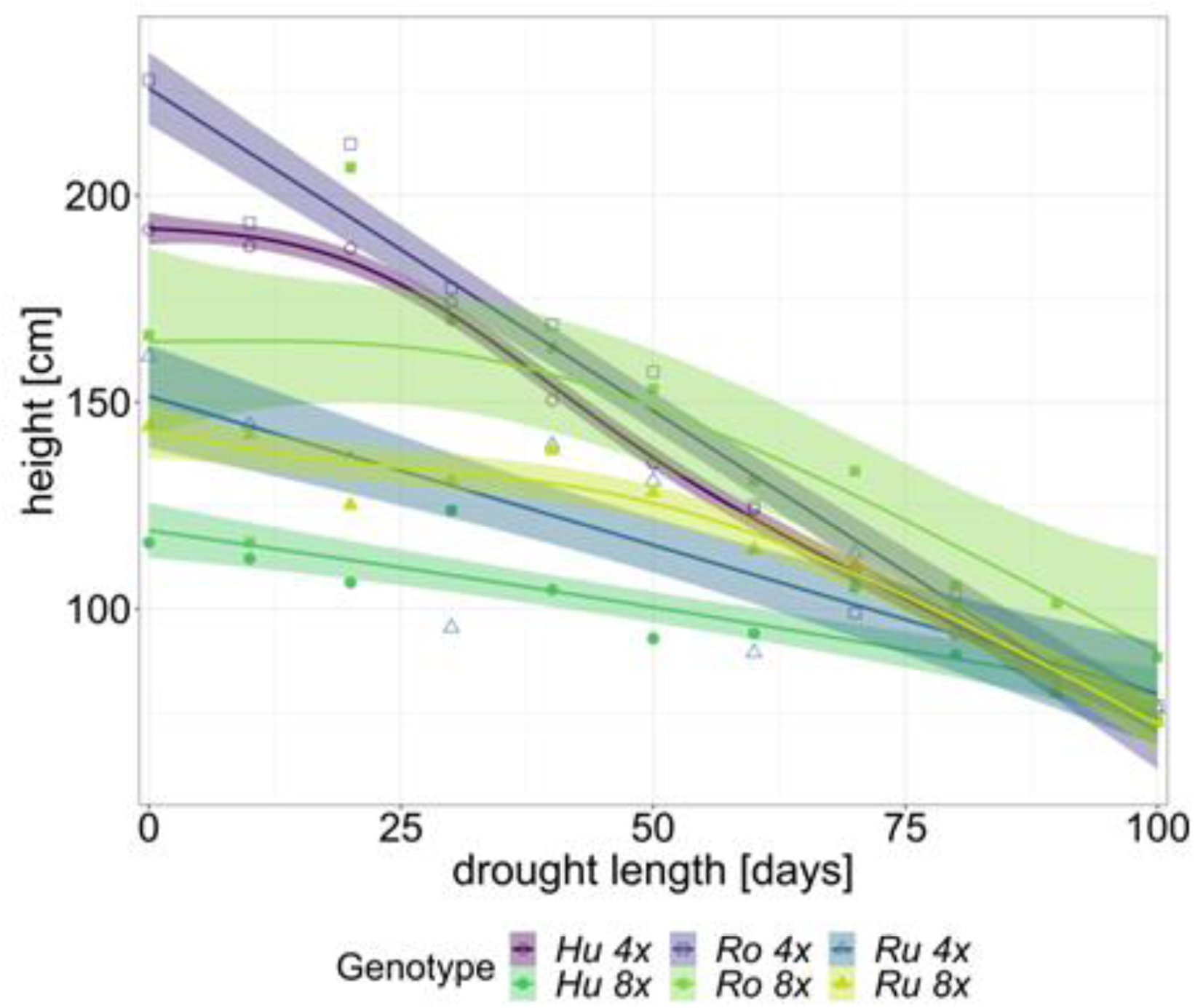
Height [cm] at harvest of the Hungarian (Hu, circles), Romanian (Ro, squares) and Russian (Ru, triangles) 4× (open symbols) and 8× (closed symbols) *Phragmites australis* genotypes at harvest along the drought length gradient. Lines show the generalized additive model; shaded areas show the 83% confidence intervals.

## Discussion

### Tetraploids outperform octoploids in absence of stress

Overall, *P. australis* reacted to drought as it has been described in previous studies: Under drought stress, the plants stopped growth and started to die off aboveground (Saltmarsh *et al*., 2006; Li *et al*., 2013; da Cunha Cruz *et al*., 2019; Huang *et al*., 2022). The longer the drought plants had to endure, the lower their biomass and the smaller the plants were at harvest (Asamoah & Bork, 2010; Touchette *et al*., 2010; Didiano *et al*., 2018; da Cunha Cruz *et al*., 2020; Huang *et al*., 2022).

However, the ploidy levels behaved contrary to our expectation: Without and up to moderate drought stress of about 40 days drainage, 4× were mostly superior to 8× in terms of biomass and morphological traits. Under long and severe drought stress, the two ploidy levels converged in their performance at a low level. Apart from a higher leaf number and lower fractions of dead stems and leaves, which could indicate a better performance and possibly higher stress tolerance of 8× compared to 4×, 8× never outperformed 4× in any part of the drought gradient. The ‘large genome constraint’ hypothesis proposed by Knight *et al*. (2005) suggests that plant species with large genomes tend to have a lower distribution and abundance in extreme environments. At the same time, a majority of studies on various species demonstrated that plants with a higher ploidy level exhibited an improved drought resistance compared to their diploid relatives (Tossi *et al*., 2022). The “original”, diploid form of *P. australis* does not exist anymore, instead tetra- and octoploids are the most common ploidy levels (Gorenflot, 1976). In the absence of stress, 8× have been shown to develop longer and thicker stems than 4× (Hanganu *et al*., 1999; Pauca-Comanescu *et al*., 1999; Hansen *et al*., 2007), while, in our study, 8× developed shorter and thinner stems than 4× without stress. In line with our results, 8× were found to have a lower stem density (stem number) and thus lower total biomass than 4× (Hanganu *et al*., 1999). Nevertheless, the opposite finding of 4× being taller and developing less stems than 8× has also been reported (Meyerson *et al*., 2016) and our results confirm other studies which conclude that a higher ploidy level does not necessarily result in bigger plants in *P. australis* (Clevering & Lissner, 1999; Hansen *et al*., 2007).

Photosynthetic activity does not provide a conclusive explanation for the better growth of 4× compared to 8× we observed in this study. The ploidy levels did not differ in their photosynthetic rate along the drought gradient. Our study did not endorse a positive effect of short drought on photosynthesis-related traits (Pagter *et al*., 2005; Saltmarsh *et al*., 2006) as the trend to limited performance started already at the shortest drought levels and became significant after 10 to 20 days of drought for biomass, morphology and photosynthetic rate, in line with the findings of Pagter *et al*. (2005) on a negative influence of mild water stress (60% watering) on leaf production and leaf growth rate. Studies reporting a positive effect of low drought stress levels used smaller plant pots and shorter drought periods (Pagter *et al*., 2005; Saltmarsh *et al*., 2006). Possibly, the treatments of 10 to 20 days drainage, which were considered a short drought in our study, already created enough water stress to have a negative effect on *P. australis*.

The intended drought length determined the timepoint of drought onset in our study. Plants that experienced long drought were drained early in their growth phase, thus stopping their development at an early stage whereas plants that experienced short drought were drained late in their growth phase. Timepoint of drought onset may have influenced mainly two aspects of our results. Firstly, the magnitude of differences between 4× and 8×: The greater height and stem number of 8× compared to 4× developed only during the course of the growing season. Plants that experienced long drought were halted in their development at a stage when 4× and 8× were still similar in height and stem number. Plants that experienced short drought, on the other hand, were drained when the differences between 4× and 8× had already developed. Secondly, biomass and morphology at harvest along the drought gradient: Plants that were drained early only had little time to grow before their growth was stopped by drought. In addition to the more severe drought stress these plants experienced, compared to their short drought counterparts, this may be an additional reason for the lower biomass and smaller plant size at harvest.

### Geographic origin can overrule ploidy effects

Previous studies did not indicate a clear trend on the reaction of different *P. australis* ploidy levels to environmental stress. In response to salt stress, for instance, different scenarios were observed: 8× being more affected by salinity than 4× (Hanganu *et al*., 1999), no effect of ploidy level on salinity tolerance (Achenbach *et al*., 2013) and salt stress lessening the differences between ploidy levels (Pauca-Comanescu *et al*., 1999), which corresponds most closely with our results. These greatly varying results could be due to the strong influence of genotype and origin on the phenotype of *P. australis* (Hansen *et al*., 2007; Achenbach *et al*., 2013) and these factors were not always excluded in previous studies on *P. australis* ploidy. A strength of our study is the examination of three genotypes per ploidy level and mitigating the influence of origin by employing a pair of 4× and 8× genotypes from each geographic origin.

In many of the examined traits, we observed a large variation between single genotypes in this study, too. Regarding biomass and morphological traits at harvest, the Hungarian 4× and Romanian 4× were often bigger (greater height, greater stem diameter) and had more stems and more biomass than the other genotypes. 4× of Hungarian and Romanian origin often outperformed their 8× counterparts while 4× and 8× of Russian origin often did not differ in these traits. This pattern was however not consistent in further traits beyond biomass and morphology, and genotypes showed a variation not clearly related to ploidy level in, for example, photosynthetic rate and belowground root mass fraction.

Climatic conditions at the genotypes’ origins could offer an explanation as to why the Russian ploidy pair behaves somewhat differently than those of the other two origins. While long-term monthly average air temperatures at the Hungarian and Romanian origins are very similar to the air temperatures during the experiment in Greifswald, the monthly average air temperatures at the Russian origin are considerably lower than in Greifswald (Figure 5). *P. australis* genotypes retain their adaptation to climatic conditions at their origin, even if they are cultivated under different climatic conditions over a longer time period (Hansen *et al*., 2007). It is therefore possible that the adaptation of the Russian genotypes to different climatic conditions as the other genotypes influences their performance (Eller *et al*., 2017). Furthermore, it is worth noting that the Hungarian and Romanian genotypes as well as the Russian 4× genotype phylogenetically belong to the mostly undifferentiated *P. australis* core group complex, while the Russian 8× genotype belongs to a group of East Asian and Australian octoploids (Lambertini *et al*., 2006). According to Clevering & Lissner (1999), the Hungarian and Romanian genotypes used in this study come from an area dominated by tetraploid *P. australis*, while the Russian genotypes come from an area dominated by octoploid *P. australis*. The location of the experiment, Greifswald, also lies in an area dominated by tetraploid *P. australis* (Clevering & Lissner, 1999). These studies thereby show other possible reasons as to why the Russian ploidy pair performs differently from the other two origin pairs. Clearly, genotypic differences beyond ploidy are important in *P. australis* and studies on the effects of ploidy should consider multiple genotypes per ploidy.

**Figure 5:**
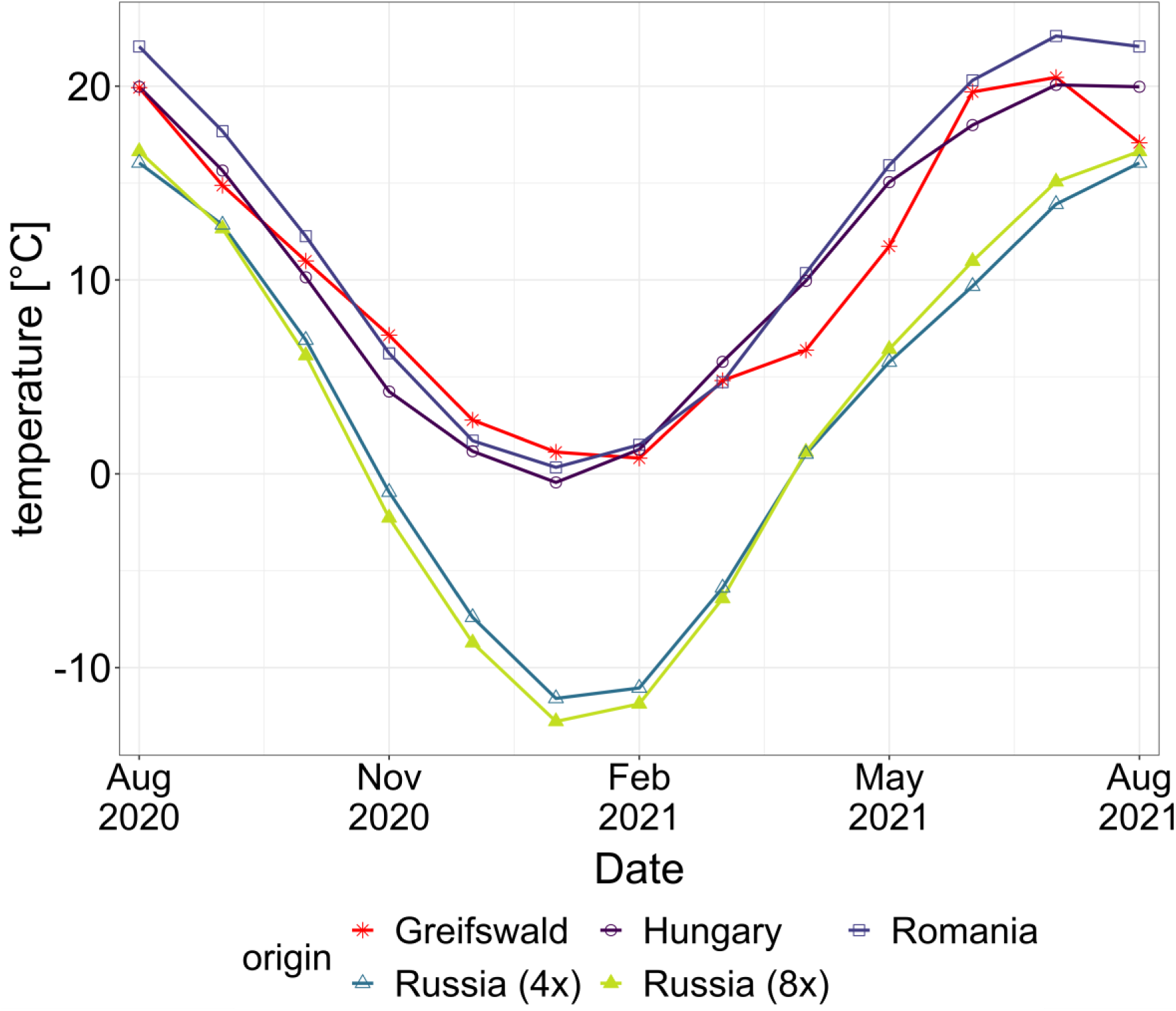
Monthly mean air temperature [°C] during the experiment in Greifswald (red stars), at the origin of the Hungarian genotypes (purple circles), at the origin of the Romanian genotypes (dark blue squares) and at the origin of the Russian genotypes (4×: blue triangles, 8×: green triangles).

Despite the strong negative effect of drought on plant performance, we assume that the plants did not die even after 100 days of drainage. While our target plants were harvested for below-ground biomass after the drought treatment finished, we observed regrowth in all but one of the plants used as buffer around the target plants within eight weeks after removing the roofs and allowing the plants to be watered by rain. Aboveground death and survival of belowground organs may be a stress avoidance strategy. Especially in established plants, rhizomes can survive and be viable even after severe drought and carbohydrates stored in rhizomes can be used as a resource for new growth (Mann *et al*., 2013; Ma *et al*., 2020).

### Implications for paludiculture in cool-temperate climates

For a profitable paludiculture with *P. australis*, drought should be avoided whenever possible. Even a short and comparatively mild drought had a negative impact on growth and biomass at harvest. This implies that in a year with drought, a reduced harvest or even complete crop failure should be expected. However, in all likelihood the plants will thrive again in the following season if sufficient water is available. Based on the studied genotypes, tetraploid *P. australis* is preferable to octoploids for cultivation under cool-temperate climatic conditions of the study region with constant water availability and mild to moderate drought (up to 40 days of drought), but under severe drought both ploidy levels show the same, very poor, performance.

## Conclusion

Drought led to the cessation of growth and subsequent dieback of aboveground biomass. Even short drought (about 20 days of drainage) had significant negative effects on biomass, morphological and physiological traits of 4× and 8× *P. australis*. In a paludiculture with *P. australis*, drought should be avoided whenever possible and a reduced harvest can be expected in a year with drought. 4× generally showed a steeper decrease in performance than 8× in the first part of the drought gradient. Under constant water availability and up to moderate drought (about 40 to 50 days of drainage), 4× outperformed 8× in almost all biomass and morphological traits, but not in photosynthetic rate. Under severe and prolonged drought, both ploidy levels performed equally poorly in our experiment. Our experiment with ploidy-pairs from different geographic origins suggests that the observed strong effect of ploidy level on plant performance under non-drought conditions is not present in all origin pairs, indicating that other effects but ploidy also play a role in plant performance and emphasizes the importance of employing genotypes of multiple origins to study ploidy effects in *P. australis*.

## Acknowledgements

We thank Aarhus University for offering the opportunity to use reed genotypes from their common garden in this study and for providing information on the genotypes; Kai-Uwe Schwarz and Heike Meyer from the Julius-Kühn Institute, Braunschweig, for invaluable effort propagating the genotypes, the gardeners of Greifswald University for much appreciated help with the plants, our student assistants for their help and all members of the Experimental Plant Ecology working group at Greifswald University for input on the experimental design, support with executing the experiment and discussions about the study.

## Author contributions

KH, KK, MS, MB and JK conceived the research and designed the experiment. JK, MS and FE provided resources for the research. KH, KK, AF and AG performed the experiment. KH, KK, CH, AF, AG and NK collected the data. KH, KK and JK analyzed the data. KH and JK wrote the original draft of the manuscript, all authors reviewed the manuscript. All authors approved the manuscript.

## Data availability

Raw data used for this manuscript will be made publicly available upon publication of the manuscript.

