## Supplemental files for "Higher ploidy coincides with inferior performance and no difference in stress tolerance in reed"

**Methods S1** Calculation of volumetric soil moisture

Internal soil moisture count of the soil moisture sensors was converted to volumetric soil moisture using the TMS Calibr tool (TOMST s.r.o., Prague, Czech Republic). Since our substrate was a peat-sand mixture, no standard calibration curve was available. We conducted an experiment to determine the calibration curve for our substrate

($y=6.81\times{10}^{-9}x^{2}+1.43\times{10}^{-4}x-0.135$) to use in the TMS Calibr tool. Two sensors measuring matrix potential (TEROS 21 Soil Water Potential Sensor, METER Group, Inc. USA, Pullman, WA, USA), one sensor measuring volumetric soil water content (Decagon 5TM Sensor, METER Group, Inc. USA, Pullman, WA, USA) and one soil moisture sensor as used in the mesocosms ( TMS-4, TOMST s.r.o., Prague, Czech Republic) were placed in a plastic tube with the same substrate used in the mesocosms; the substrate was then dried out in a climate cabinet until the permanent wilting point of pF 4.2 was reached. Durig the drying process, the matrix potential was check regularly by manual measurements with a field tensiometer (Tensio 100, Umwelt-Geräte-Technik GmbH, Müncheberg, Germany). A quadratic model was developed from the measured values of the volumetric soil water content ($y$) and the internal device count of the TMS-4 ($x$).

**Methods S2** Summary of 2020 experiment

In 2020, plants were subjected to a gradient of drought length with 11 treatment levels, ranging from zero to 40 days. Starting on 6th August 2020, water was drained from a container every 4 days, one container remained as a control without drainage.

Plant growth (maximum height, number of alive stems) was measured bi-weekly from 11th June to 17th September, photosynthetic rate was measured on three days before harvest of aboveground biomass on 17th September 2020. During the drought treatment, soil moisture decreased to values between 22.1 % (pF 2.2) and 12.8 % (pF 2.8) which are well above the permanent wilting point of our substrate (pF 4.2 = 5.2 % soil moisture), indicating that only a mild drought was reached. Nevertheless, plants stayed small in the experiment. At harvest, 4x reached a maximum height of 66 cm (0 days drought treatment), a maximum number of 22 stems (0 days drought treatment), a mean stem diameter of 1.7 mm (0 days drought treatment) and an aboveground dry weight of 4.2 g (0 days drought treatment). 8x reached a maximum height of 62 cm (40 days drought treatment), a maximum number of 17 stems (0 days drought treatment), a mean stem diameter of 2.0 mm (40 days drought treatment) and an aboveground dry weight of 4.1 g (40 days drought treatment). In 2020, drought only caused a significant increase of aboveground biomass at harvest of 8x, but had no significant effect on biomass of 4x and on height, number of stems and mean stem diameter at harvest as well as photosynthetic rate of both ploidy levels. 4x produced significantly more aboveground biomass (0-10 days drought) and greater stem number at harvest (0-30 days drought) than 8x, but 8x had a significantly higher mean stem diameter than 4x (15-40 days drought) and there was no significant difference between ploidy levels regarding height at harvest and photosynthetic rate.

After harvest, all containers were filled with tap water up to soil surface again. Plants were fertilized with 8 g of fertilizer (Hakaphos Blau, 15 N + 10 P + 15 K + 2 Mg + trace elements, Compo Expert, Münster, Germany) per tube on 24th September and 13th November. During winter, from 20th November 2020 to 11th April 2021, water was removed from the containers and the containers were wrapped in bubble wrap to protect the plants from the cold.

On 14th May 2021, the last measurement before the start of the treatments in 2021, a positive effect of the 2020 treatment on plant growth could be observed as height of 8x increased significantly with increasing 2020 drought length while height of 4x and number of living stems of both ploidy levels showed no significant effect of the 2020 treatment (height: gam: adjusted R2 = 0.743, 4x: edf = 1.155, F-value = 0.026, p <1, 8x: edf = 1.0, F-value = 20.991, p<0.001; stem: gam: adjusted R2 = 0.472, 4x: edf = 1.248, F-value = 0.106, p<1, 8x: edf = 1.0, F-value = 0.0, p<1).

**Methods S3** Calculation of biomass fractions

Aboveground mass fraction was calculated as allocation of aboveground biomass relative to total plant biomass.

Belowground root mass fraction was calculated as allocation of root biomass relative to total belowground biomass, consisting of root and rhizome biomass.

As aboveground biomass was divided into an alive and dead fraction at harvest, fraction of dead aboveground biomass was calculated relative to total aboveground biomass; fraction of dead stems in total number of stems at harvest was calculated as well as fraction of dead leaves per stem relative to total number of leaves per stem.

**Notes S1** Development of stem number under drought treatments

Onset of drought led to a decrease in the number of alive stems in both 4x and 8x (Figure S1). Only in the control treatment, there was no significant decline in the number of living stems of both 4x and 8x from the highest stem number to the end of the treatment. Already in the treatment with 10 days drainage number of living stems of 8x decreased significantly by 13% from the highest stem number to the end of the treatment, and in treatments with longer drainage number of living stems decreased significantly by 18%, 64%, 80%, 74%, 93%, 98% in the treatments with 20,30,40,50,60 and 70 days drainage, respectively, and by 100% in the treatments with 80 days and longer drainage. A decrease in number of living stems of 4x, while not significant in the 10 days drainage treatment, was significant in the treatments with 20 days and longer drainage, decreasing by 13%, 74%, 98%, 92% and 99% in the treatments with 20,30,40,50 and 70 days of drainage and by 100% in the treatments with 80, 90 and 100 days of drainage. 4x developed significantly more living stems than 8x over the course of the growing season in most treatments (control, 10, 20, 40, 50, 70, 80 and 100 days drainage, Table 2). Onset of drought mostly resulted in numbers of living stems of the two ploidy levels converging as stem number decreased.

**Notes S2** Effect of drought length on biomass allocation

Fraction of aboveground biomass in total biomass (aboveground mass fraction, Figure S2c, Table 2) decreased significantly with increasing length of drought in both ploidy levels. 4x had a higher aboveground mass fraction than 8x over a large part of the gradient (about 30 – 100 days drought). Fraction of roots in belowground biomass (belowground root mass fraction, Figure S2d, Table 2) increased significantly and linearly with increasing length of drought in both ploidy levels. Under 0 to about 60 days drought, 4x had a significantly lower belowground root mass faction (and consequently a significantly higher belowground rhizome mass fraction) than 8x, under prolonged drought the ploidy levels did not differ.

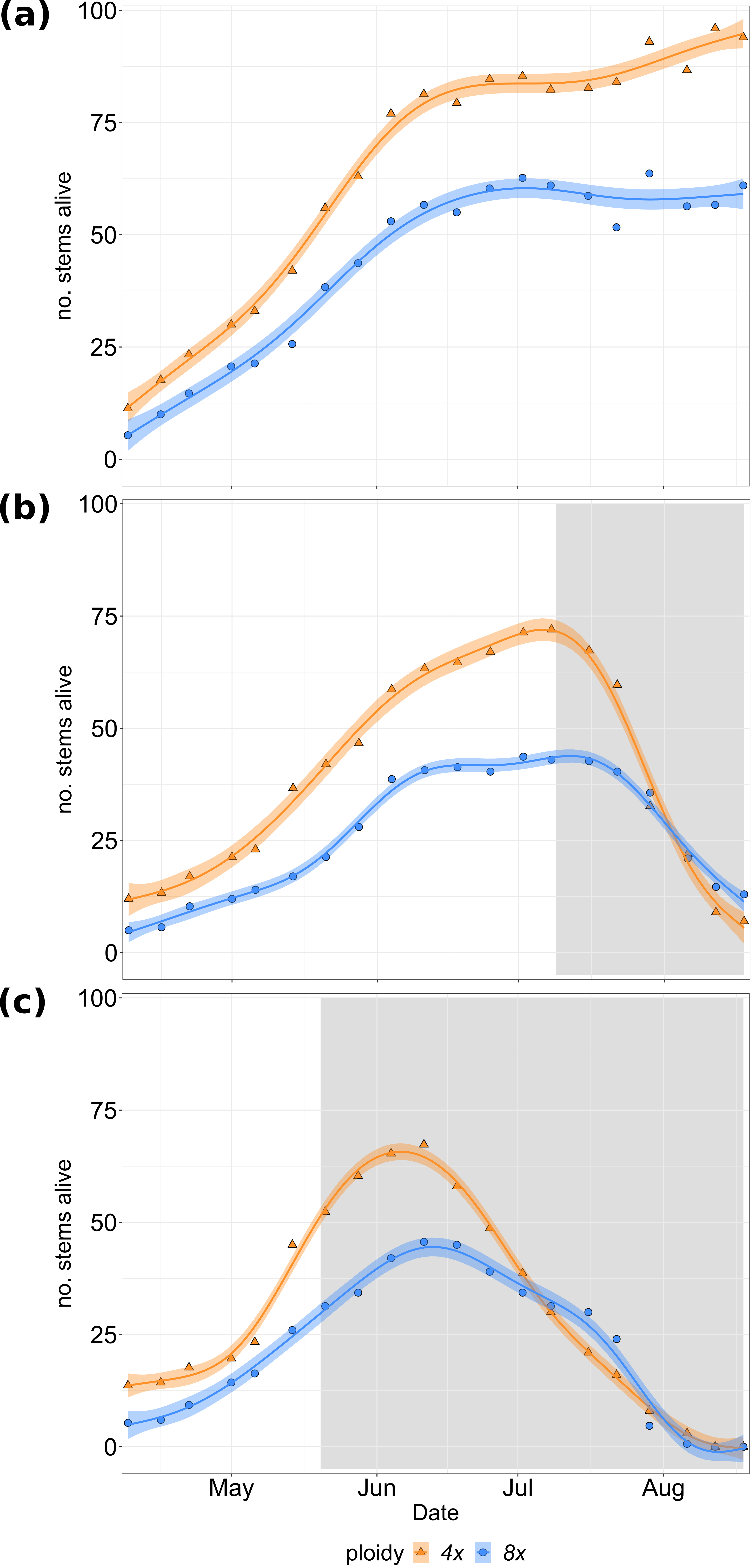
**Figure S1** Number of alive stems during the growing season 2021 of 4x (orange, triangles) and 8x (blue, circles) *Phragmites australis* genotypes in the treatments (a) no drought, (b) 50 days drainage and (c) 100 days drainage. Lines show the generalized additive model, shaded areas show the 83% confidence intervals. Gray shading in the background depicts the timeperiod of drainage in the respective treatment.

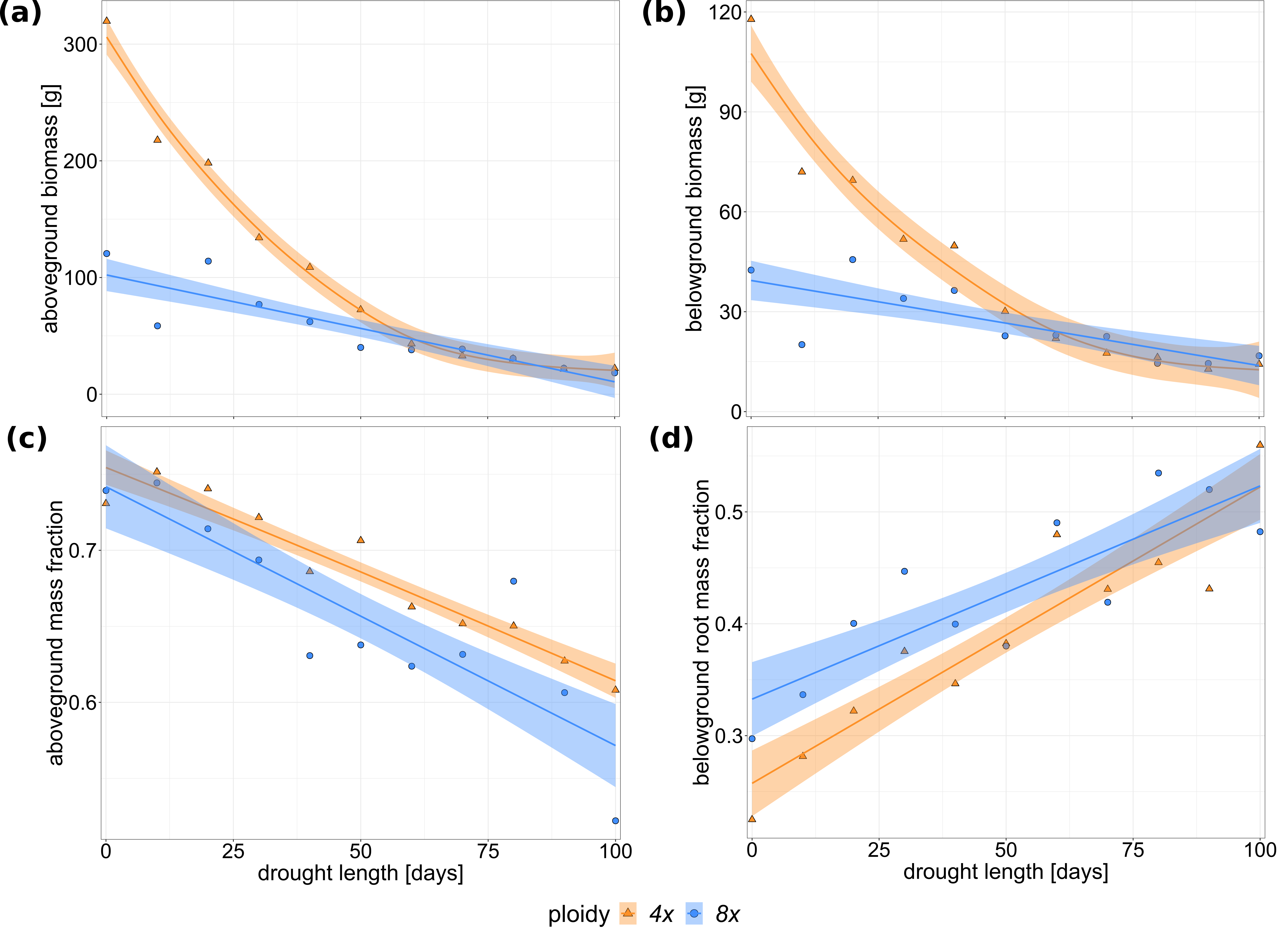
**Figure S2** (a) Aboveground biomass [g], (b) belowground biomass [g], (c) fraction of aboveground biomass in total biomass (aboveground mass fraction) and (d) fraction of root biomass in belowground biomass (belowground root mass fraction) of 4x (orange, triangles) and 8x (blue, circles) *Phragmites australis* genotypes along the drought length gradient. Lines show the generalized additive model ((a) aboveground biomass: adjusted R^2^ = 0.957, 4x: edf = 3.51, F-value = 87, p < 0.001, 8x: edf = 1, F-value = 36.34, p < 0.001; (b) belowground biomass: adjusted R^2^ = 0.913, 4x: edf = 3.182, F-value = 46.63, p < 0.001; 8x: edf = 1, F-value = 12.66, p < 0.01; (c) aboveground mass fraction: adjusted R^2^ = 0.929, 4x: edf = 1, F-value = 90.62, p < 0.001, 8x: edf = 5.073, F-value = 29.72, p < 0.001; belowground root mass fraction: adjusted R^2^ = 0.778, 4x: edf = 1, F-value = 47.30, p < 0.001, 8x: edf = 1, F-value = 24.47, p < 0.001). Shaded areas show the 83% confidence intervals.

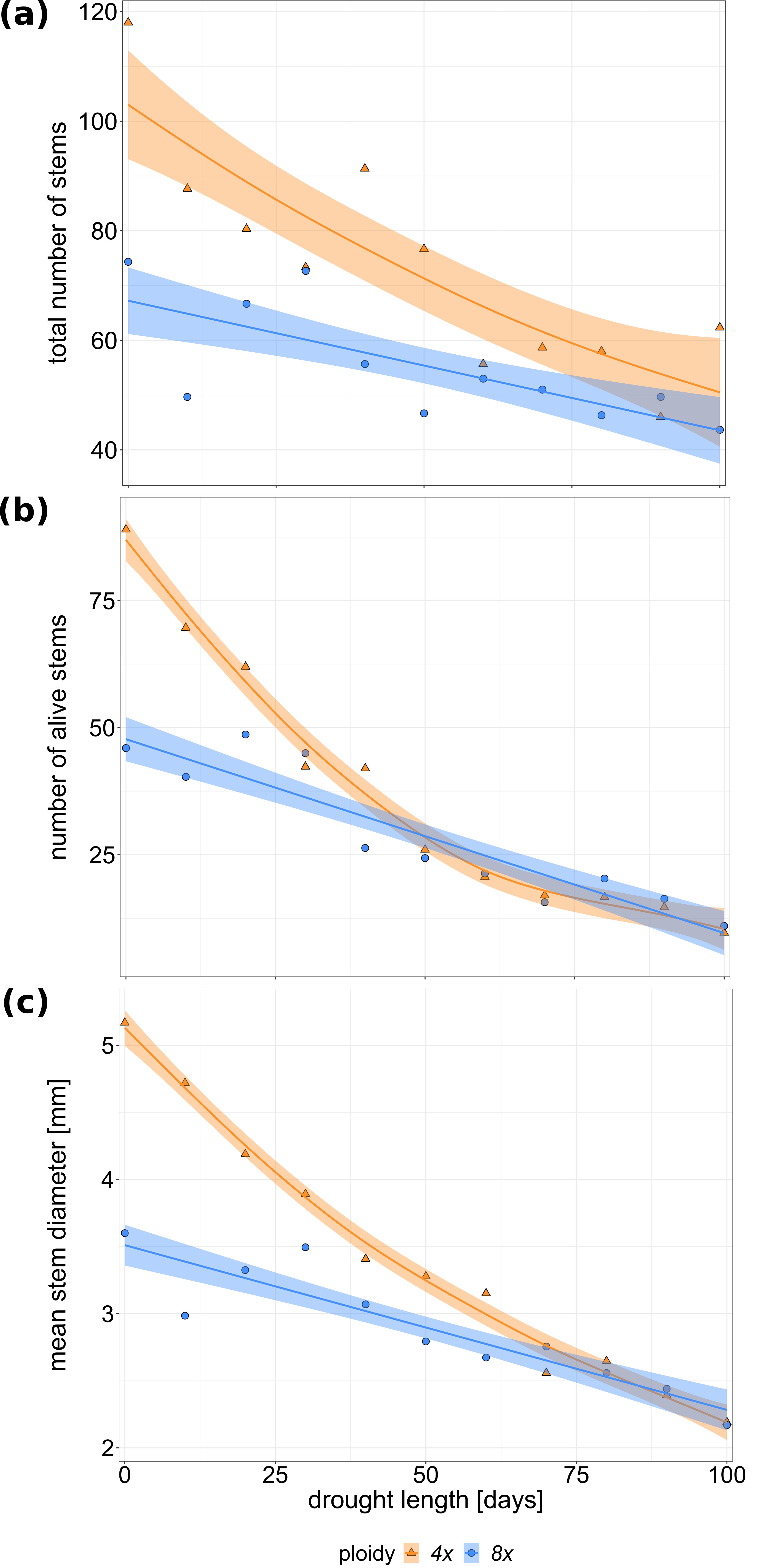
**Figure S3** (a) Total number of stems, (b) number of alive stems and (c) mean stem diameter [mm] of 4x (orange, triangles) and 8x (blue, circles) *Phragmites australis* genotypes at harvest along the drought length gradient. Lines show the generalized additive model ((a) total number of stems: adjusted R^2^ = 0.745, 4x: edf = 1.916, F-value = 15.187, p < 0.001, 8x: edf = 1, F-value = 7.056, p < 0.05; (b) number of alive stems: adjusted R^2^ = 0.958, 4x: edf = 3.222, F-value = 92.49, p < 0.001, 8x: edf = 3.045, F-value = 24.40, p < 0.001; (c) mean stem diameter: adjusted R^2^ = 0.956, 4x: edf = 2.527, F-value = 108.59, p < 0.001, 8x: edf = 1, F-value = 60.17, p < 0.001). Shaded areas show the 83% confidence intervals.

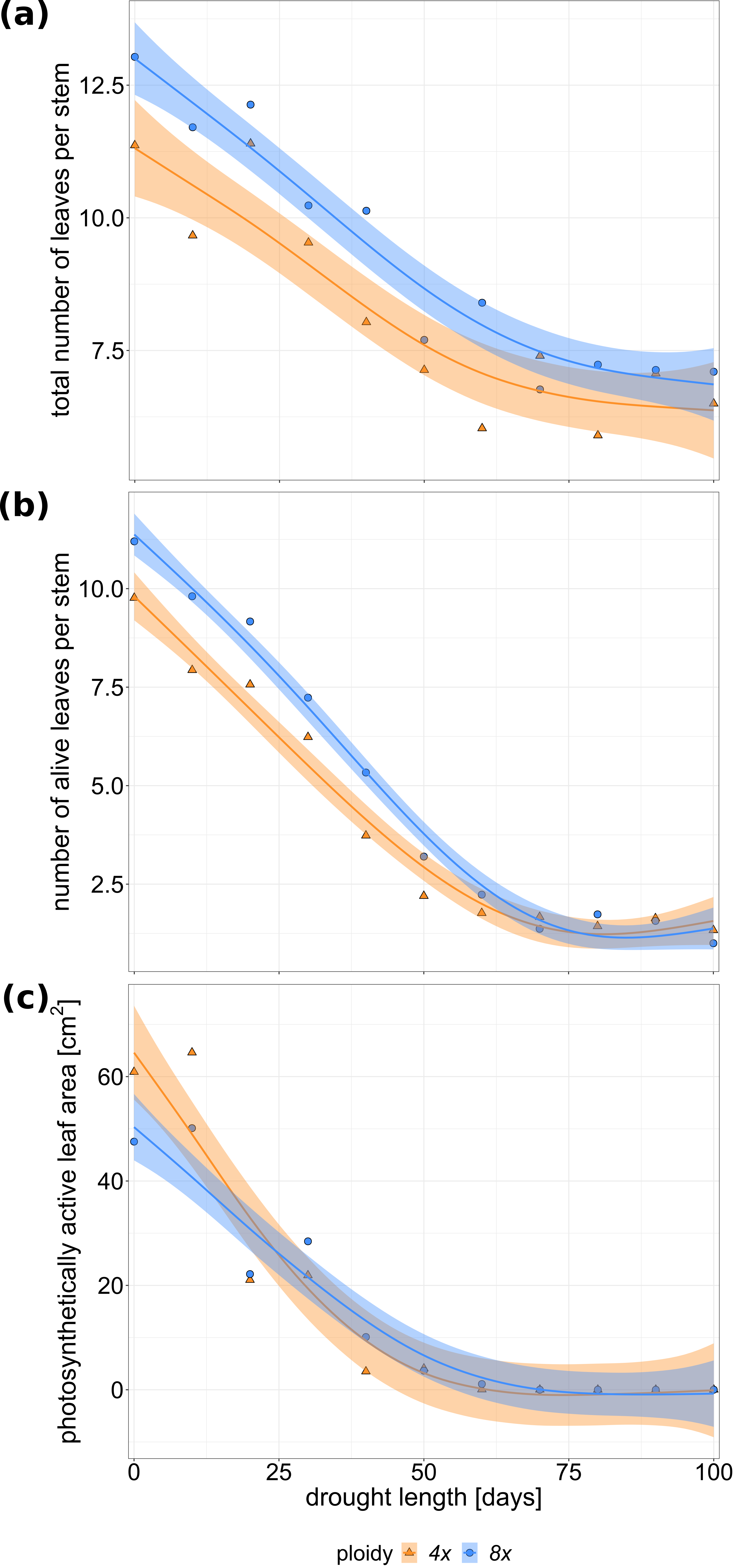
**Figure S4** (a) Total number of leaves per stem, (b) number of alive leaves per stem and (c) photosynthetically active leaf area [cm^2^] of 4x (orange, triangles) and 8x (blue, circles) *Phragmites australis* genotypes at harvest along the drought length gradient. Number of (alive) leaves per stem was counted on the 10 longest stems of each genotype per level. Lines show the generalized additive model ((a) total number of leaves per stem: adjusted R^2^ = 0.878, 4x: edf = 2.609, F-value = 18.05, p < 0.001, 8x: edf = 2.278, F-value = 29.60, p < 0.001; (b) number of alive leaves per stem: adjusted R^2^ = 0.979, 4x: edf = 2.776, F-value = 129.8, p < 0.001, 8x: edf = 2.829, F-value = 192.4, p < 0.001; (c) photosynthetically active leaf area: adjusted R^2^ = 0.897, 4x: edf = 3.246, F-value = 28.3, p < 0.001, 8x: edf = 2.568, F-value = 22.5, p < 0.001). Shaded areas show the 83% confidence intervals.

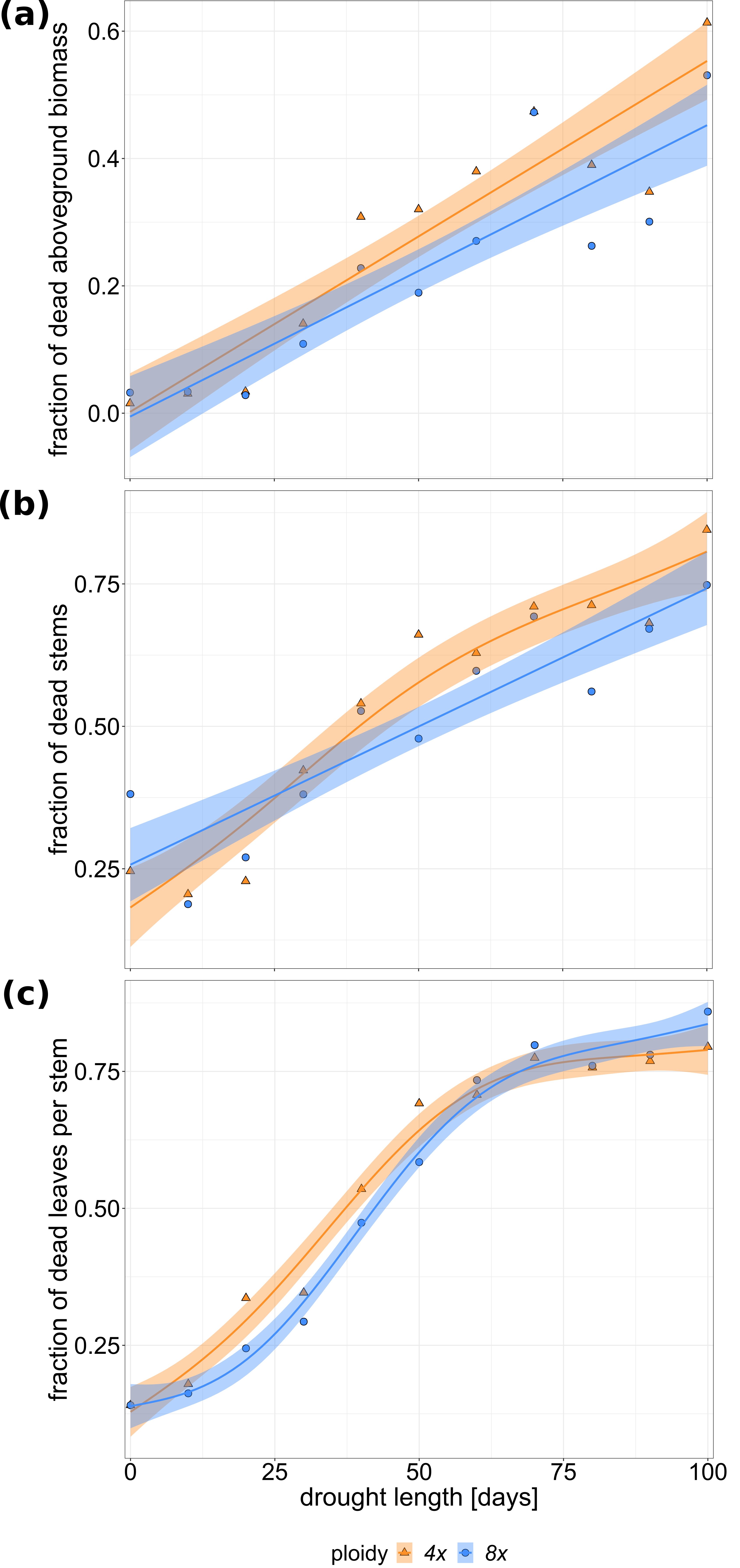
**Figure S5** (a) Fraction of dead aboveground biomass in total aboveground biomass, (b) fraction of dead in total number of stems and (c) fraction of dead leaves in total number of leaves per stem of 4x (orange, triangles) and 8x (blue, circles) *Phragmites australis* genotypes at harvest along the drought length gradient. Number of leaves per stem was counted on the 10 longest stems of each genotype per level. Lines show the generalized additive model ((a) fraction of dead aboveground biomass: adjusted R^2^ = 0.805, 4x: edf = 1, F-value = 51.69, p < 0.001; 8x: edf = 1, F-value = 35.64, p < 0.001; (b) fraction of dead stems: adjusted R-squared = 0.85, 4x: edf = 1.884, F-value = 32.68, p < 0.001; 8x: edf = 1; F-value = 43.56, p < 0.001; (c) fraction of dead leaves per stem: adjusted R-squared = 0.981, 4x: edf = 3.421, F-value = 127.4, p < 0.001; 8x: edf = 3.599, F-value = 150.4, p < 0.001). Shaded areas show the 83% confidence intervals.

**Table S1** Analysis of communal tap water used in the mesocosm experiment. Tap water was a mixture of water from waterworks Schönwalde (60%) and Hohenmühl (40%). Values in this table are calculated accordingly from data of Stadtwerke Greifswald GmbH (n.d.).

| Parameter | Unit | Method | Value |
| --- | --- | --- | --- |
| pH |  | DIN EN ISO 10523 | 7.39 |
| **NH_4_^+^** | [mg L^-1^] | DIN EN ISO 11732 | 0.014 |
| **NO_2_^-^** | [mg L^-1^] | DIN EN ISO 13395 | 0.016 |
| **NO_3_^2-^** | [mg L^-1^] | DIN EN ISO 10304-1 | 1.72 |
| **PO_4_^3-^** | [mg L^-1^] | DIN EN ISO 15681-1 | 0.038 |
| **K^+^** | [mg L^-1^] | DIN EN ISO 11885 | 2.84 |

**Table S2** Generalized additive model used for each dependent variable. Results of each model summary are presented in the figure captions.

| **Dependent variable** | **Generalized additive model** |
| --- | --- |
| **Biomass at harvest** | |
| Total biomass | gam(total biomass ~ ploidy + s(drought length, by = ploidy, bs = “tp”), method = “REML”) |
| Aboveground biomass | gam(aboveground biomass ~ ploidy + s(drought length, by = ploidy, bs = “tp”), method = “REML”) |
| Belowground biomass | gam(belowground biomass ~ ploidy + s(drought length, by = ploidy, bs = “tp”), method = “REML”) |
| **Morphology at harvest** | |
| Stem length | gam(stem length ~ ploidy + s(drought length, by = ploidy, bs = “tp”), method = “REML”) |
| Total number of stems | gam(total number of stems ~ ploidy + s(drought length, by = ploidy, bs = “tp”), method = “REML”) |
| Alive number of stems | gam(alive number of stems ~ ploidy + s(drought length, by = ploidy, bs = “tp”), method = “REML”) |
| Mean stem diameter | gam(mean stem diameter ~ ploidy + s(drought length, by = ploidy, bs = “tp”), method = “REML”) |
| Total leaves per stem | gam(total number of leaves per stem ~ ploidy + s(drought length, by = ploidy, bs = “tp”), method = “REML”) |
| Alive leaves per stem | gam(number of alive leaves per stem ~ ploidy + s(drought length, by = ploidy, bs = “tp”, k = 4), method = “REML”) |
| **Dead biomass fractions** | |
| Fraction of dead aboveground biomass in total aboveground biomass | gam((aboveground dead biomass / aboveground biomass) ~ ploidy + s(drought length, by = ploidy, bs = “tp”), method = “REML”) |
| Fraction of dead in total number of stems | gam((number of dead stems / total number of stems) ~ ploidy + s(drought length, by = ploidy, bs = “tp”, k = 9), method = “REML”) |
| Fraction of dead leaves in total number of leaves per stem | gam(number of alive leaves per stem ~ ploidy + s(drought length, by = ploidy, bs = “tp”, k = 5), method = “REML”) |
| **Biomass fractions** | |
| Aboveground mass fraction | gam((aboveground biomass / total biomass) ~ ploidy + s(drought length, by = ploidy, bs = “tp”, k = 7), method = “REML”) |
| Belowground root mass fraction | gam((root biomass / belowground biomass) ~ ploidy + s(drought length, by = ploidy, bs = “tp”), method = “REML”) |
| **Photosynthetic rate** | |
| Photosynthetic rate | gam(photosynthetic rate ~ ploidy + s(drought length, by = ploidy, bs = “tp”, k = 3), method = “REML”) |
| Photosynthetically active leaf area | gam(photosynthetically active leaf area ~ ploidy + s(drought length, by = ploidy, bs = “tp”), method = “REML”) |

**Table S3** Results of the model check of each generalized additive model used (Table S1). Model quality was checked using gam.check (package mgcv).

| **Dependent variable** | **Console output** | **Residual plots** |
| --- | --- | --- |
| **Biomass at harvest** | |  |
| Total biomass | Method: REML; Optimizer: outer newton. Full convergence after 10 iterations. \| k’ \| edf \| k-index \| p-value; s(drought length):Ploidy 4x \| 9.00 \| 3.47 \| 0.8 \| 0.12; s(drought length):Ploidy 8x \| 9.00 \| 1.00 \| 0.8 \| 0.11 | 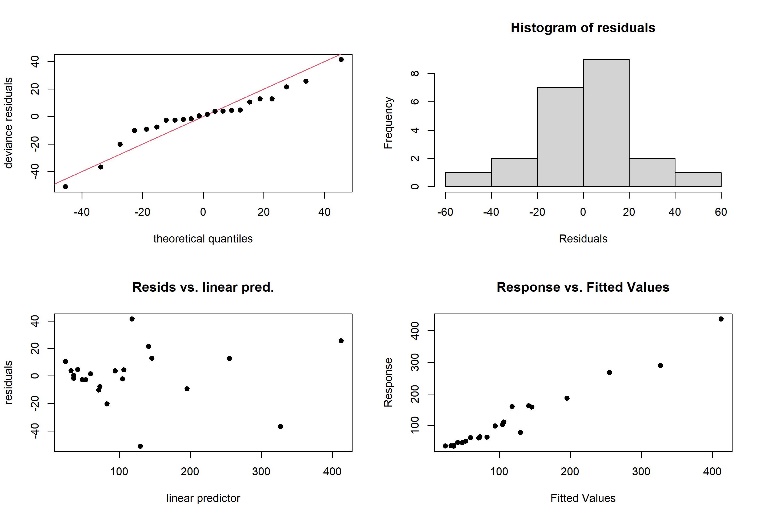 |
| Aboveground biomass | Method: REML; Optimizer: outer newton. Full convergence after 10 iterations. \| k’ \| edf \| k-index \| p-value; s(drought length):Ploidy 4x \| 9.00 \| 3.51 \| 0.85 \| 0.19; s(drought length):Ploidy 8x \| 9.00 \| 1.00 \| 0.85 \| 0.20 | 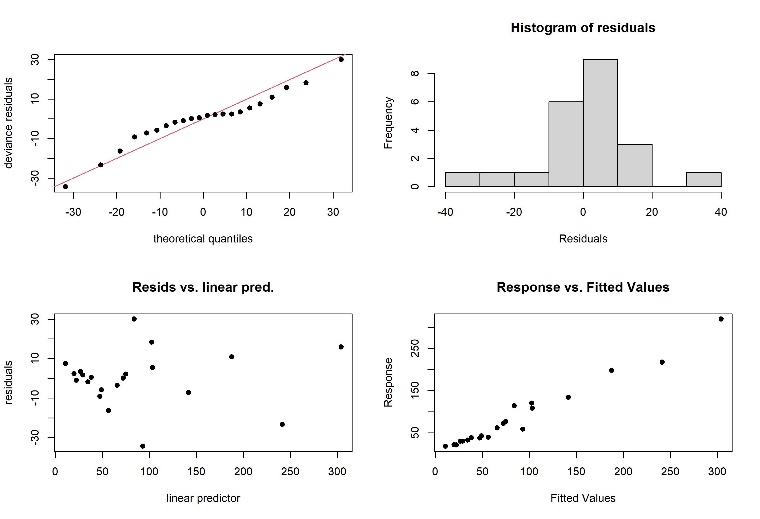 |
| Belowground biomass | Method: REML; Optimizer: outer newton. Full convergence after 10 iterations. \| k’ \| edf \| k-index \| p-value; s(drought length):Ploidy 4x \| 9.00 \| 3.18 \| 0.68 \| 0.07 .; s(drought length):Ploidy 8x \| 9.00 \| 1.00 \| 0.68 \| 0.03 * | 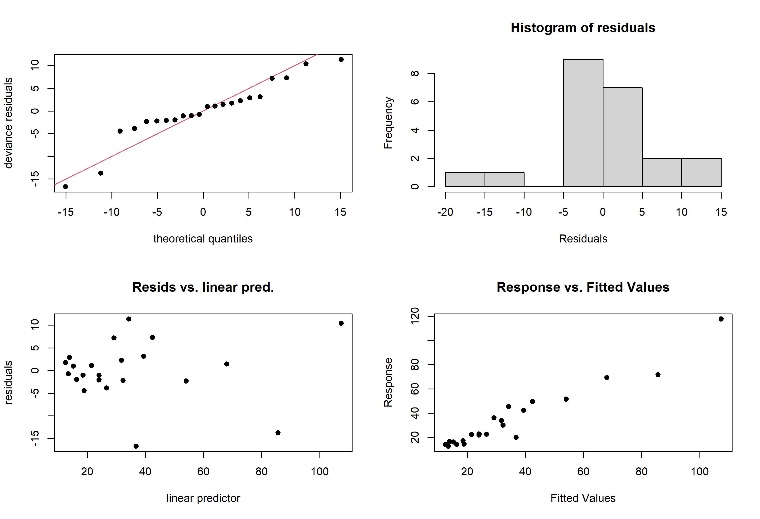 |
| **Morphology at harvest** | |  |
| Stem length | Method: REML; Optimizer: outer newton. Full convergence after 10 iterations. \| k’ \| edf \| k-index \| p-value; s(drought length):Ploidy 4x \| 9.00 \| 1.00 \| 1.02 \| 0.43; s(drought length):Ploidy 8x \| 9.00 \| 2.69 \| 1.02 \| 0.42 | 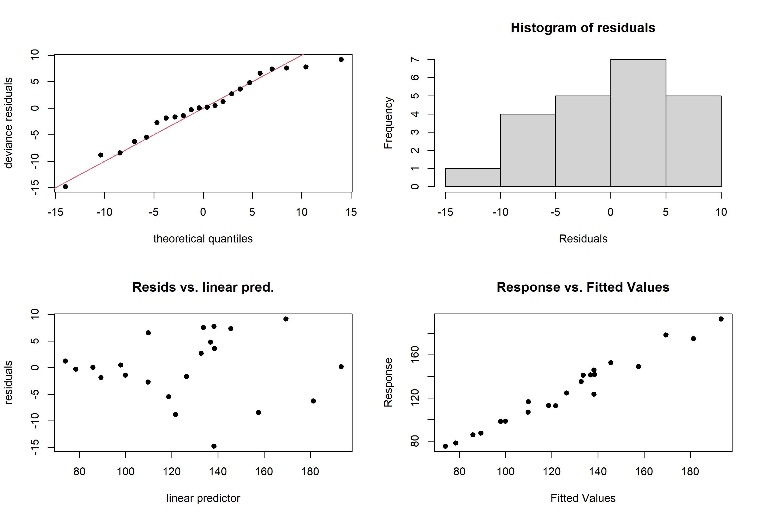 |
| Total number of stems | Method: REML; Optimizer: outer newton. Full convergence after 8 iterations. \| k’ \| edf \| k-index \| p-value; s(drought length):Ploidy 4x \| 9.00 \| 1.92 \| 0.77 \| 0.10; s(drought length):Ploidy 8x \| 9.00 \| 1.00 \| 0.77 \| 0.11 | 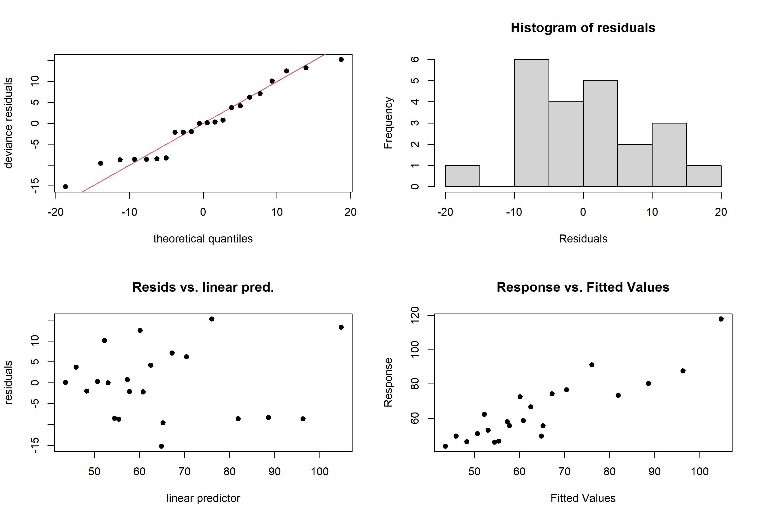 |
| Alive number of stems | Method: REML; Optimizer: outer newton. Full convergence after 5 iterations. \| k’ \| edf \| k-index \| p-value; s(drought length):Ploidy 4x \| 9.00 \| 3.22 \| 1.05 \| 0.48; s(drought length):Ploidy 8x \| 9.00 \| 3.04 \| 1.05 \| 0.54 | 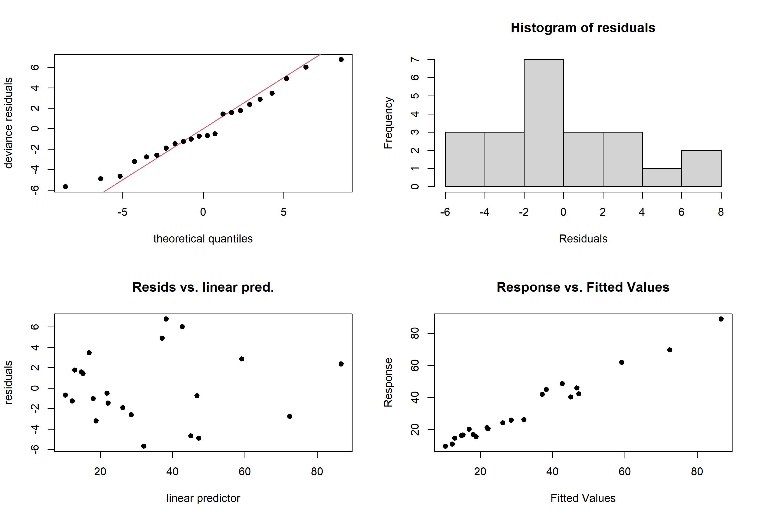 |
| Mean stem diameter | Method: REML; Optimizer: outer newton. Full convergence after 12 iterations. \| k’ \| edf \| k-index \| p-value; s(drought length):Ploidy 4x \| 9.00 \| 2.53 \| 1.16 \| 0.68; s(drought length):Ploidy 8x \| 9.00 \| 1.00 \| 1.16 \| 0.72 | 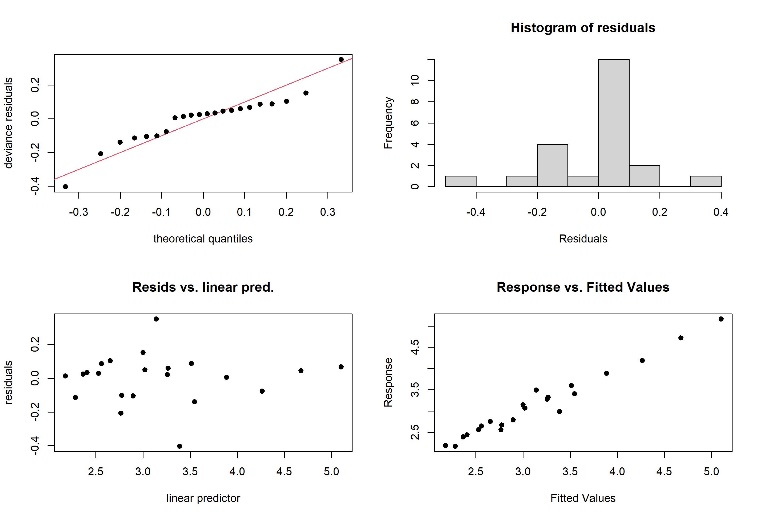 |
| Total leaves per stem | Method: REML; Optimizer: outer newton. Full convergence after 4 iterations. \| k’ \| edf \| k-index \| p-value; s(drought length):Ploidy 4x \| 9.00 \| 2.61 \| 0.78 \| 0.095 .; s(drought length):Ploidy 8x \| 9.00 \| 2.28 \| 0.78 \| 0.130 | 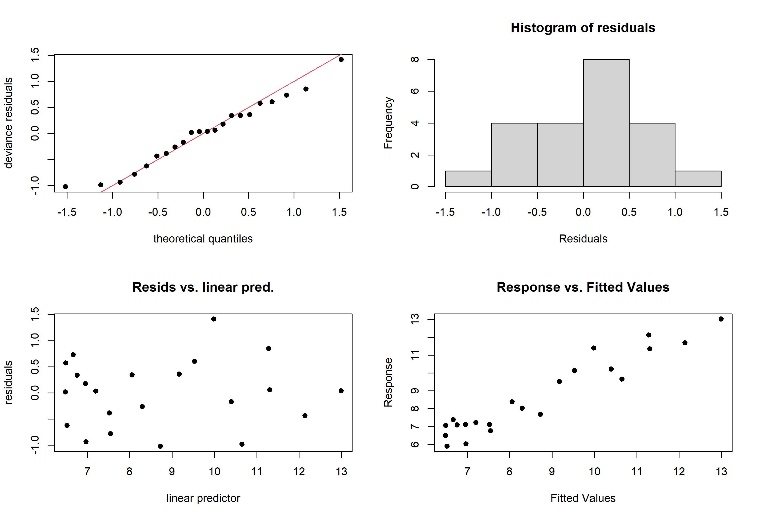 |
| Alive leaves per stem | Method: REML; Optimizer: outer newton. Full convergence after 7 iterations. \| k’ \| edf \| k-index \| p-value; s(drought length):Ploidy 4x \| 3.00 \| 2.78 \| 0.51 \| 0.01 **; s(drought length):Ploidy 8x \| 3.00 \| 2.83 \| 0.51 \| < 2e^-16^ *** | 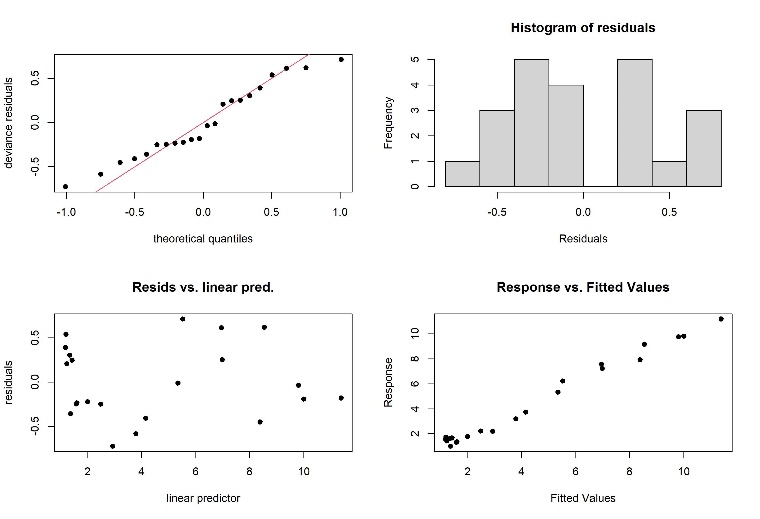 |
| **Dead biomass fractions** | |  |
| Fraction of dead aboveground biomass in total aboveground biomass | Method: REML; Optimizer: outer newton. Full convergence after 12 iterations. \| k’ \| edf \| k-index \| p-value; s(drought length):Ploidy 4x \| 9 \| 1 \| 0.59 \| 0.015 *; s(drought length):Ploidy 8x \| 9 \| 1 \| 0.59 \| 0.020 * | 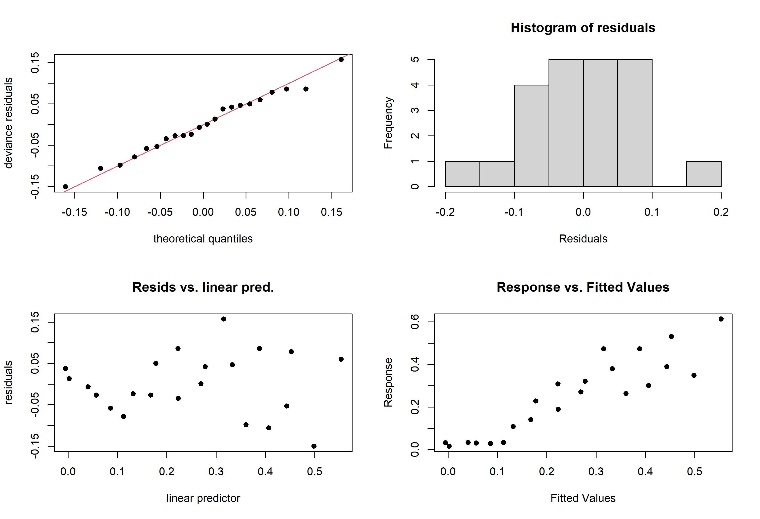 |
| Fraction of dead in total number of stems | Method: REML; Optimizer: outer newton. Full convergence after 10 iterations. \| k’ \| edf \| k-index \| p-value; s(drought length):Ploidy 4x \| 8.00 \| 1.88 \| 0.52 \| 0.005 **; s(drought length):Ploidy 8x \| 8.00 \| 1.00 \| 0.52 \| < 2e^-16^ *** | 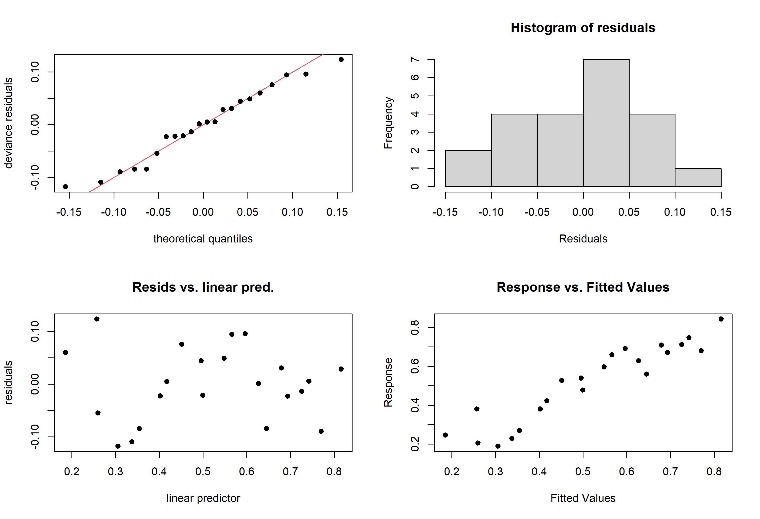 |
| Fraction of dead leaves in total number of leaves per stem | Method: REML; Optimizer: outer newton. Full convergence after 10 iterations. \| k’ \| edf \| k-index \| p-value; s(drought length):Ploidy 4x \| 4.00 \| 3.42 \| 0.85 \| 0.14; s(drought length):Ploidy 8x \| 4.00 \| 3.60 \| 0.85 \| 0.15 | 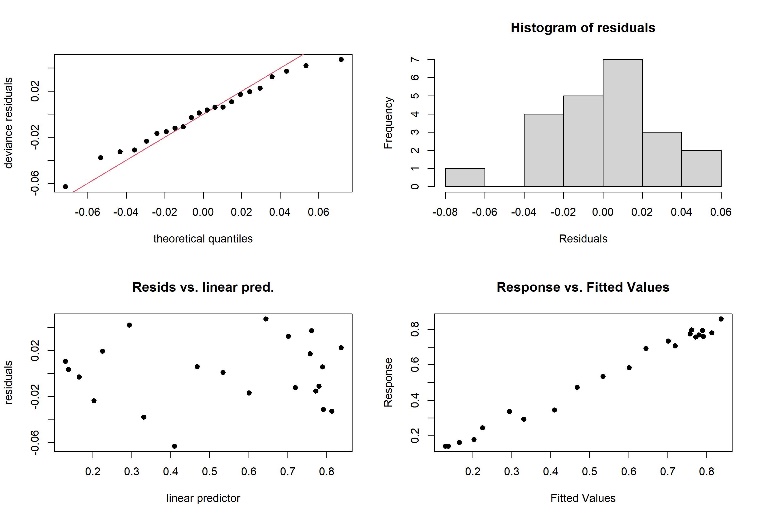 |
| **Biomass fractions** | |  |
| Aboveground mass fraction | Method: REML; Optimizer: outer newton. Full convergence after 7 iterations. \| k’ \| edf \| k-index \| p-value; s(drought length):Ploidy 4x \| 6.00 \| 1.00 \| 0.86 \| 0.23; s(drought length):Ploidy 8x \| 6.00 \| 5.07 \| 0.86 \| 0.15 | 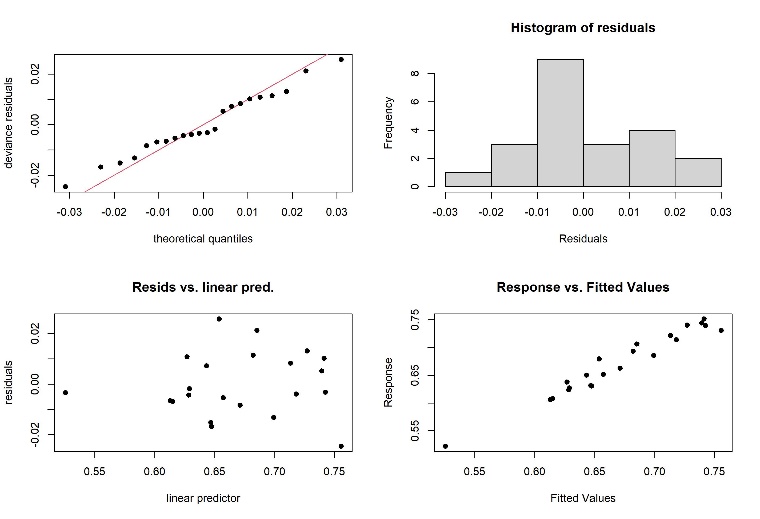 |
| Belowground mass fraction | Method: REML; Optimizer: outer newton. Full convergence after 7 iterations. \| k’ \| edf \| k-index \| p-value; s(drought length):Ploidy 4x \| 6.00 \| 1.00 \| 0.86 \| 0.22; s(drought length):Ploidy 8x \| 6.00 \| 5.07 \| 0.86 \| 0.16 | 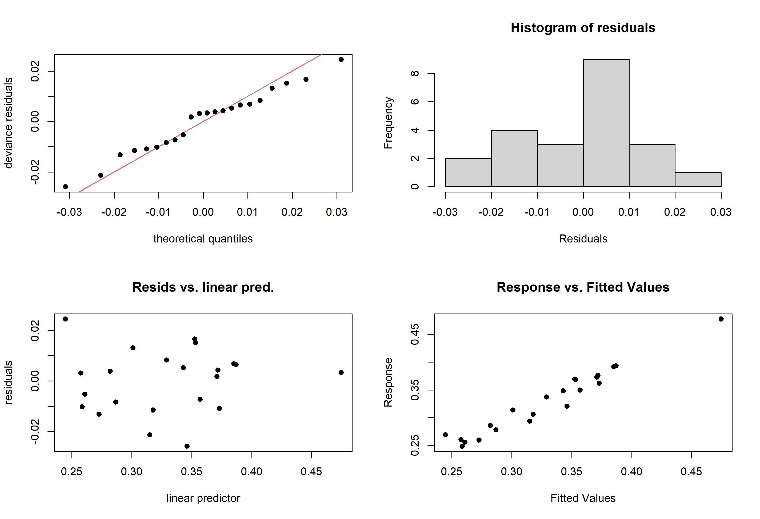 |
| Belowground root mass fraction | Method: REML; Optimizer: outer newton. Full convergence after 10 iterations. \| k’ \| edf \| k-index \| p-value; s(drought length):Ploidy 4x \| 9 \| 1 \| 1.04 \| 0.52; s(drought length):Ploidy 8x \| 9 \| 1 \| 1.04 \| 0.44 | 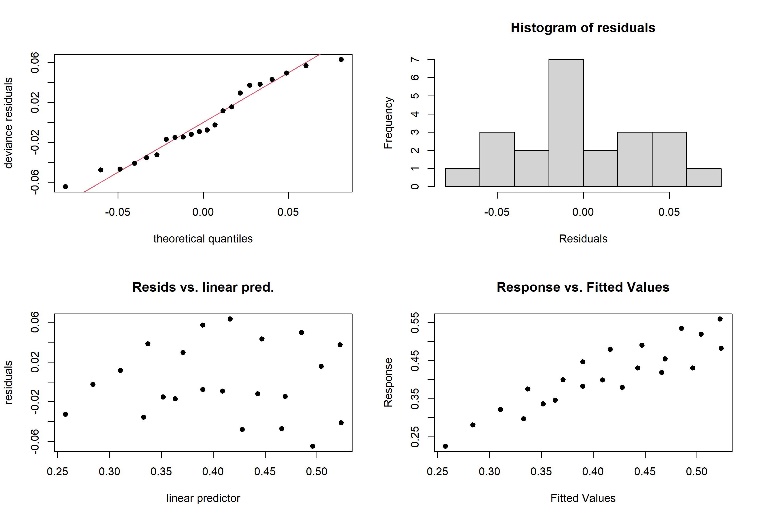 |
| Belowground rhizome mass fraction | Method: REML; Optimizer: outer newton. Full convergence after 10 iterations. \| k’ \| edf \| k-index \| p-value; s(drought length):Ploidy 4x \| 9 \| 1 \| 1.04 \| 0.46; s(drought length):Ploidy 8x \| 9 \| 1 \| 1.04 \| 0.36 | 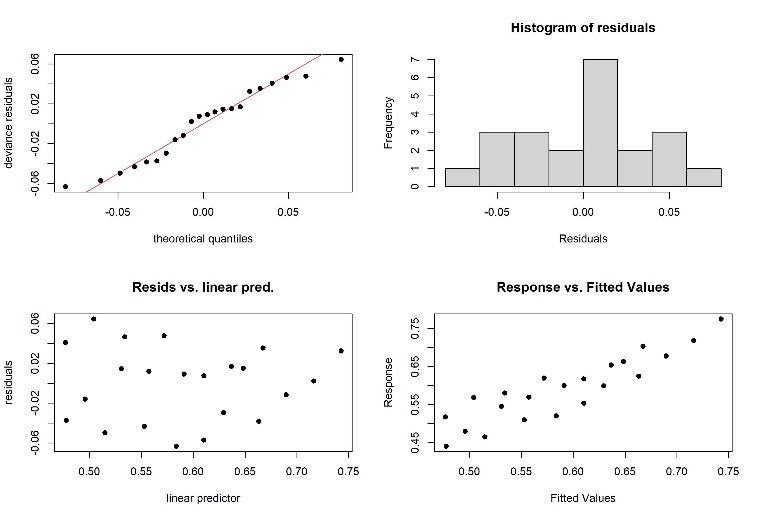 |
| **Photosynthetic rate** | |  |
| Photosynthetic rate | Method: REML; Optimizer: outer newton. Full convergence after 4 iterations. \| k’ \| edf \| k-index \| p-value; s(drought length):Ploidy 4x \| 2.00 \| 1.59 \| 1.18 \| 0.64; s(drought length):Ploidy 8x \| 2.00 \| 1.42 \| 1.18 \| 0.62 | 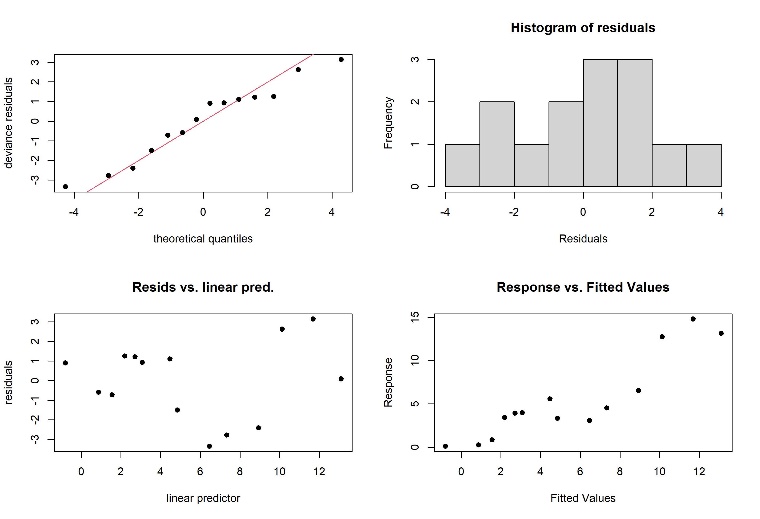 |
| Photosynthetically active leaf area | Method: REML; Optimizer: outer newton. Full convergence after 3 iterations. \| k’ \| edf \| k-index \| p-value; s(drought length):Ploidy 4x \| 9.00 \| 3.25 \| 0.92 \| 0.23; s(drought length):Ploidy 8x \| 9.00 \| 2.57 \| 0.92 \| 0.32 | 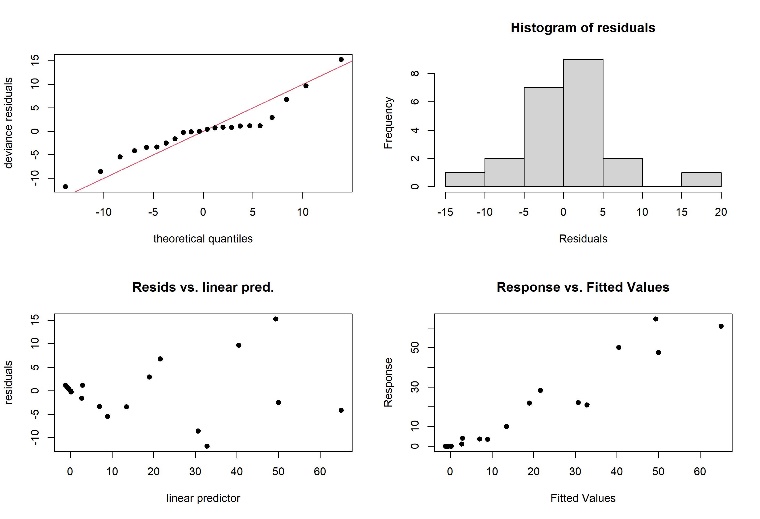 |
| **Aboveground biomass element content** | |  |
| Total N content | Method: REML; Optimizer: outer newton. Full convergence after 5 iterations. \| k’ \| edf \| k-index \| p-value; s(drought length):Ploidy 4x \| 9.00 \| 2.47 \| 0.95 \| 0.29; s(drought length):Ploidy 8x \| 9.00 \| 2.73 \| 0.95 \| 0.26 | 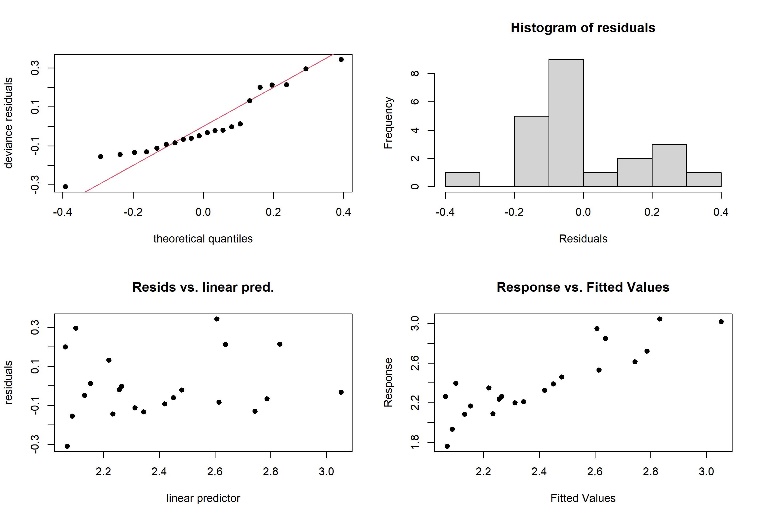 |
| P content | Method: REML; Optimizer: outer newton. Full convergence after 11 iterations. \| k’ \| edf \| k-index \| p-value; s(drought length):Ploidy 4x \| 2.00 \| 1.86 \| 0.95 \| 0.3; s(drought length):Ploidy 8x \| 2.00 \| 1.00 \| 0.95 \| 0.3 | 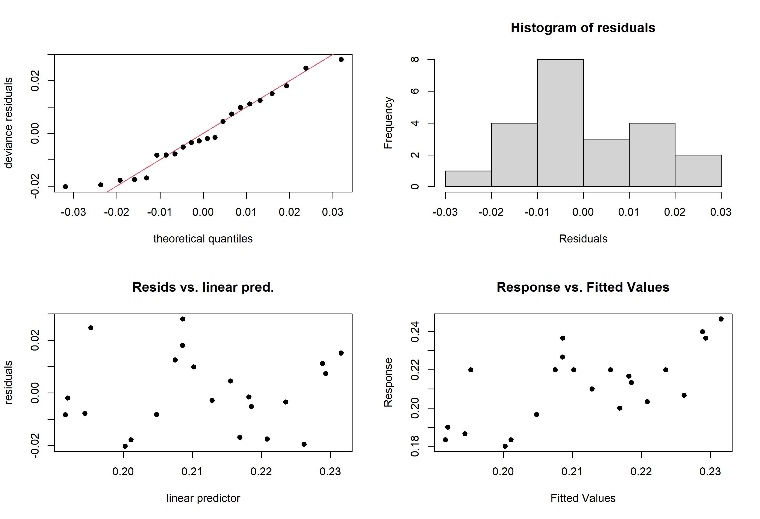 |
